# Bacterial diet modulates tamoxifen-induced death via host fatty acid metabolism

**DOI:** 10.1101/2021.11.05.467497

**Authors:** Cédric Diot, Aurian P. García-González, Andre F. Vieira, Melissa Walker, Megan Honeywell, Hailey Doyle, Olga Ponomarova, Yomari Rivera, Huimin Na, Hefei Zhang, Michael Lee, Carissa P. Olsen, Albertha J.M. Walhout

**Affiliations:** Department of Systems Biology, University of Massachusetts Chan Medical School, Worcester, MA, 01605, USA; Department of Chemistry and Biochemistry, Worcester Polytechnic Institute, 100 Institute Road, Worcester, MA, 01609, USA

## Abstract

Tamoxifen is a selective estrogen receptor (ER) modulator that is used to treat ER-positive breast cancer, but that at high doses kills both ER-positive and ER-negative breast cancer cells. We recapitulate this off-target effect in *Caenorhabditis elegans*, which does not have an ER ortholog. We find that different bacteria dramatically modulate tamoxifen toxicity in *C. elegans*, with a three-order of magnitude difference between animals fed *Escherichia coli*, *Comamonas aquatica*, and *Bacillus subtilis*. Remarkably, host fatty acid (FA) biosynthesis mitigates tamoxifen toxicity, and different bacteria provide the animal with different FAs, resulting in distinct FA profiles. Surprisingly these bacteria modulate tamoxifen toxicity by different death mechanisms, some of which are modulated by FA supplementation and others by antioxidants. Together, this work reveals a complex interplay between microbiota, FA metabolism and tamoxifen toxicity that may provide a blueprint for similar studies in more complex mammals.

## Introduction

The bacteria that inhabit our body, known as our microbiota, influence many biological processes in both health and disease, and greatly contribute to our metabolic capacity ^1, 2^. The gut microbiota are among the first cells to encounter orally ingested nutrients and xenobiotics. Therefore, it is perhaps not surprising that the microbiota can also impact our response to medications^1^. Several mechanisms have been identified by which microbes influence the host’s drug response. First, the microbial composition can change in response to drugs. Known as dysbiosis, such altered composition can affect the host’s response to therapeutic drugs^3, 4^. Second, bacteria can alter host drug availability, for instance by drug sequestration or modification^1, 5^. Third, the metabolic crosstalk between the microbiota and its host can modify host physiology, and consequently drug efficacy^6^. All these processes can change drug action and it is therefore important to systematically identify which bacteria modulate the efficacies of which drugs, and to dissect the mechanisms involved.

Systematic identification of microbial effects on the host’s drug response in humans or mammalian model systems is challenging because of their complex diet and microbiota, as well as relatively low scalability. Recently, we and others, have developed the nematode *Caenorhabditis elegans* and its bacterial diet as an interspecies model system to rapidly assess the animal’s response to different drugs and how this response can be modulated by different bacteria^7–9^. The application of high-throughput genetic screens not only in the host, but also in the bacteria it consumes, make this a powerful model^4, 10–12^. Recent studies illustrate the insights on host-bacteria-drug interactions that can be obtained using this model system. In a screen with 11 chemotherapeutic drugs and two bacterial diets, we found that three drugs, 5-fluoro-2’-deoxyuridine (FUDR), 5-fluoro-uracil (5-FU) and camptothecin (CPT) elicited a reproductive phenotype, two of which were modulated by two bacterial species, in opposite directions ^7^. Specifically, a diet of *Escherichia coli* rendered the animals more sensitive to FUDR, but less sensitive to CPT, relative to a diet of *Comamonas aquatica*^7^. We found that bacterial pyrimidine metabolism, specifically the generation of 5-fluorouridine monophosphate (FUMP), was critical to FUDR toxicity in the animal^7, 8^.

Tamoxifen is a triphenylethylene belonging to the class of drugs known as selective estrogen receptor modulators (SERMs). SERMs bind to and either activate or repress the estrogen receptor (ER), a nuclear hormone receptor (NHR) transcription factor^13^. Tamoxifen is commonly used to treat ER-positive breast cancers, as well as other estrogen-dependent ailments such as ovarian or endometrial carcinoma ^14, 15^. A significant proportion of ER-negative breast cancers also appear responsive to tamoxifen, but the underlying biology remains to be uncovered^16^. There has also been increasing clinical interest in the potential of tamoxifen for treating various other cancer types^17^. The potential repurposing of tamoxifen for the treatment of non-ER-positive breast malignancies requires an understanding of the potential alternative mechanisms of action of this drug, especially in ER-negative models.

The *C. elegans* genome encodes more than 250 NHRs^18^. Most of these are paralogs of HNF4α and there is no ER ortholog. We find that tamoxifen is toxic in *C. elegans* at higher doses and that three bacterial species, *E. coli, C. aquatica,* and *Bacillus subtilis,* confer dramatically different levels of sensitivity to tamoxifen toxicity, with an EC_50_ ranging three orders of magnitude. We identify a role for host fatty acid (FA) metabolism in modulating tamoxifen toxicity. We then find that the FA composition of *C. elegans* is greatly modulated by bacterial diet, and that FA supplementation modulates drug toxicity in a diet-specific manner. Antioxidant supplementation and expression profiling further point to a role of lipid oxidative stress. Perhaps most strikingly, we find that different bacteria modulate *C. elegans* tamoxifen toxicity through different death mechanisms: on *E. coli* and *C. aquatica*, but not on *B. subtilis,* apoptosis deficient *ced-3* and *ced-4* mutants potentiate drug toxicity. Taken together, we show that a complex interplay between FA metabolism, oxidative stress, and differential potentials to engage cell-death pathways, modulate tamoxifen toxicity.

## Results

### Tamoxifen kills both ER-positive and ER-negative cancer cells

We first confirmed that tamoxifen can kill ER-negative breast cancer cells^19^. Specifically, we compared the effects of tamoxifen on ER-positive T-47D cells, which rely on ER for proliferation, to the effects of tamoxifen on ER-negative MDA-MB-231 cells^20^. Drug toxicity can be assessed through two parameters: growth rate (GR) and fractional viability (FV). The GR index is calculated by the relative number of live cells over time in presence or absence of drug and integrates both cytostatic and lethal effects of a drug^21^. In contrast, FV only considers the lethal effect of a drug^22^. We first calculated the GR index and, as expected, observed a decrease in the growth rate of T-47D cells incubated with tamoxifen in a dose-dependent manner (**Fig. 1a**). While lower doses of tamoxifen did not affect the growth rate of MDA-MB-231 cells, we found that concentrations greater than 10 µM resulted in a reduction in the GR, indicating either a reduced proliferation rate or an increased death rate (**Fig. 1a**). When we computed FV, we found that at tamoxifen doses greater than 10 µM, killed both T-47D and MDA-MB- 231 cells effectively (**Fig. 1b**). This indicates that tamoxifen toxicity relies on two distinct mechanisms depending on the dose: at low doses, tamoxifen slows the growth of ER-positive, but not ER-negative cells, whereas at high doses, tamoxifen kills independently of whether the cells express ER. Since this cell death does not depend on ER, it can be considered an off-target effect.

**Fig. 1:**
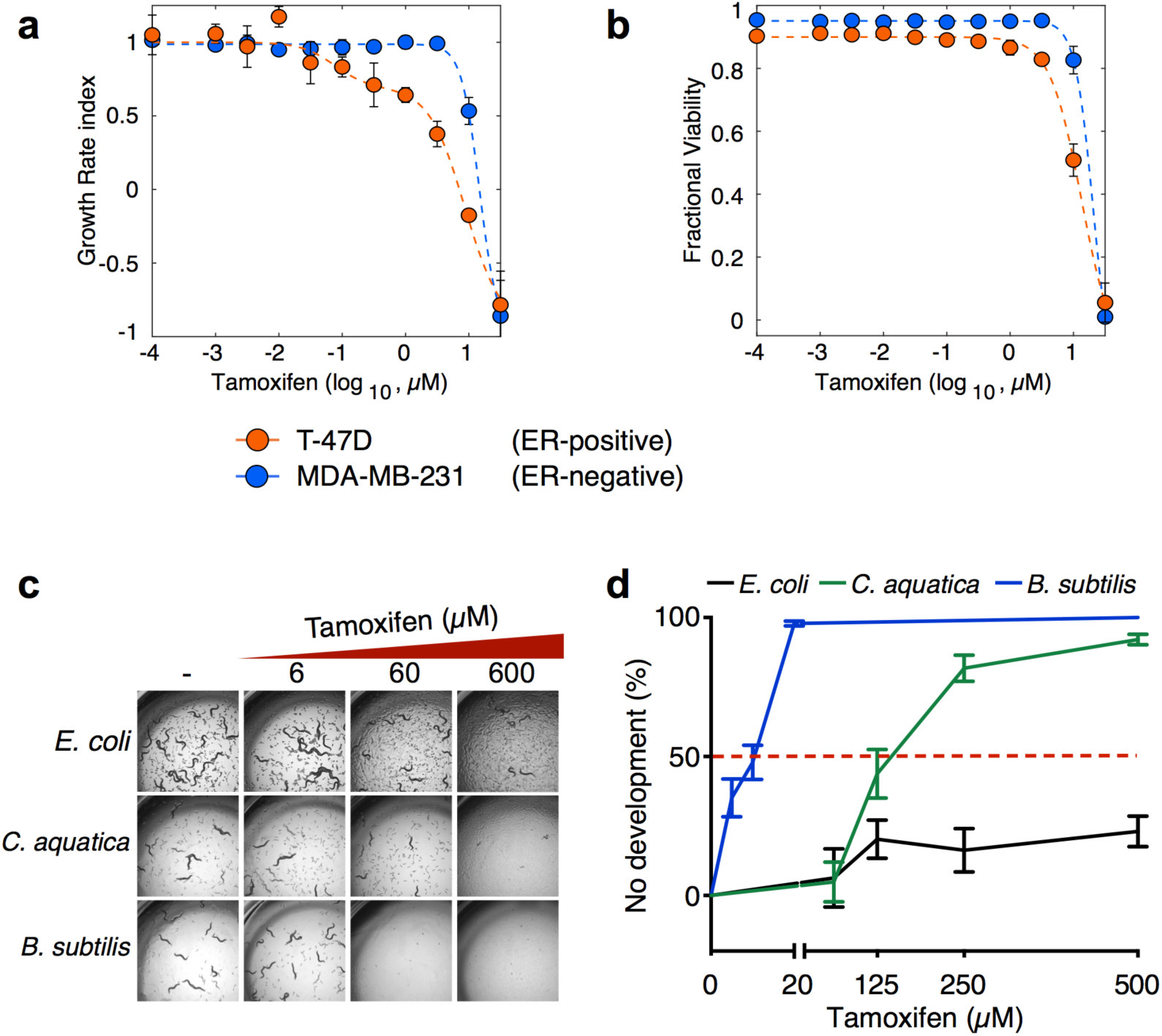
Tamoxifen Kills ER-Negative Breast Cancer Cells and Animals. **a**, **b** The growth rate index (**a**) and fractional viability (**b**) of ER-positive (T-47D) and ER-negative (MDA-MB-231) breast cancer cells plotted as a function of increasing concentrations of tamoxifen. **c** Bright-field images showing *C. elegans* supplemented with increasing doses of tamoxifen, fed *E. coli*, *C. aquatica*, and *B. subtilis*. Images were taken at 2x magnification after 48h exposure to tamoxifen. **d** Dose response curves (DRCs) of *C. elegans* supplemented with increasing doses of tamoxifen, fed *E. coli*, *C. aquatica*, and *B. subtilis.* Data are represented as mean ±SEM of three biological replicates.

### Bacteria differentially modulate tamoxifen toxicity in *C. elegans*

*C. elegans* does not have an obvious ER ortholog^18, 23^. Therefore, we wondered whether we could use *C. elegans* to study off-target tamoxifen toxicity in an intact animal. Since we previously found that the bacterial diet consumed by *C. elegans* can greatly affect the response to chemotherapeutic drugs^7^, we supplemented increasing doses of tamoxifen to larval stage 1 (L1)-arrested animals fed either of three different bacterial diets: *E. coli, C. aquatica* or *B. subtilis*. Visual inspection showed dramatic differences in tamoxifen toxicity, depending on bacterial diet: there was little effect on animal development on *E. coli*, while animals fed *C. aquatica* displayed developmental arrest or delay at high drug concentrations, and animals fed *B. subtilis* were exquisitely sensitive to tamoxifen, even at low micromolar concentrations (**Fig. 1c** and **Supplementary Fig. 1**). We next used L1 arrest as a proxy for tamoxifen toxicity and plotted the proportion of animals that failed to develop after incubation on tamoxifen-containing plates for 48 hours as a function of the drug concentration. The resulting dose response curves (DRCs) confirmed that tamoxifen is more than three orders of magnitude more toxic to animals fed *B. subtilis* than to animals fed *E. coli* (**Fig. 1d**). Notably, bacterial lawns on NGM plates were not affected by the presence of tamoxifen (**Supplementary Fig. 2**). Together, these results show that tamoxifen is toxic to *C. elegans* at high doses and that bacterial diet greatly modulates this toxicity.

### Bacteria modulate drug bioavailability

To gain insight into the mechanism by which bacteria modulate tamoxifen toxicity in *C. elegans*, we performed genetic screens in both *E. coli and C. aquatica*. Specifically, we used the *E. coli* Keio mutant collection, which contains 3,985 single-gene deletion mutant strains^24^, and a *C. aquatica* mutant collection of 5,760 strains that we previously generated by transposon-based mutagenesis^11^ (**Fig. 2a**). To enable the identification of bacterial mutants that either increase or decrease tamoxifen toxicity in *C. elegans*, we screened each bacterial mutant collection with two doses of tamoxifen: a toxic dose (200 μM and 300 μM on animals fed *C. aquatica* and *E. coli,* respectively), and a non-toxic dose (100 µM) on which animals develop on either diet. We did not identify any *E. coli* mutants that reproducibly altered drug toxicity in *C. elegans*. This indicates that active *E. coli* metabolism is unlikely to play a dominant role in modifying tamoxifen toxicity in the animal. However, we did find four *C. aquatica* mutants that changed the animal’s response to tamoxifen, two of which decreased and two that increased toxicity (**Fig. 2b**). The two mutants that increased the severity of tamoxifen toxicity harbor the transposon insertion in the *tadC* gene, which encodes a component of the type II secretion system^25^, and in the *exbD* gene, which encodes a protein involved in TonB-dependent transport^26^. The two mutants that decreased tamoxifen toxicity harbor the transposon in a gene encoding a hypothetical protein with no annotated functional domains and in the *acrR* gene, which encodes a transcriptional repressor of the *acrA/B* multidrug efflux pump^27^(**Fig. 2c**). This pump is associated with efflux of hydrophobic xenobiotics^28^. Since we identified a transcriptional repressor of this pump, this may suggest that the pump removes tamoxifen from the bacteria. These results suggest that tamoxifen transport, rather than active bacterial metabolism, affects the amount of drug taken up by *C. aquatica,* and, therefore, drug bioavailability in *C. elegans* fed this bacterial diet. To test this, we measured tamoxifen accumulation in wild-type and mutant strains of *C. aquatica* exposed to the drug by gas chromatography - mass spectrometry (GC-MS). We found that the four *C. aquatica* mutants harbor different levels of tamoxifen, and that these levels correlate with tamoxifen toxicity in *C. elegans* (**Fig. 2d**).

**Fig. 2:**
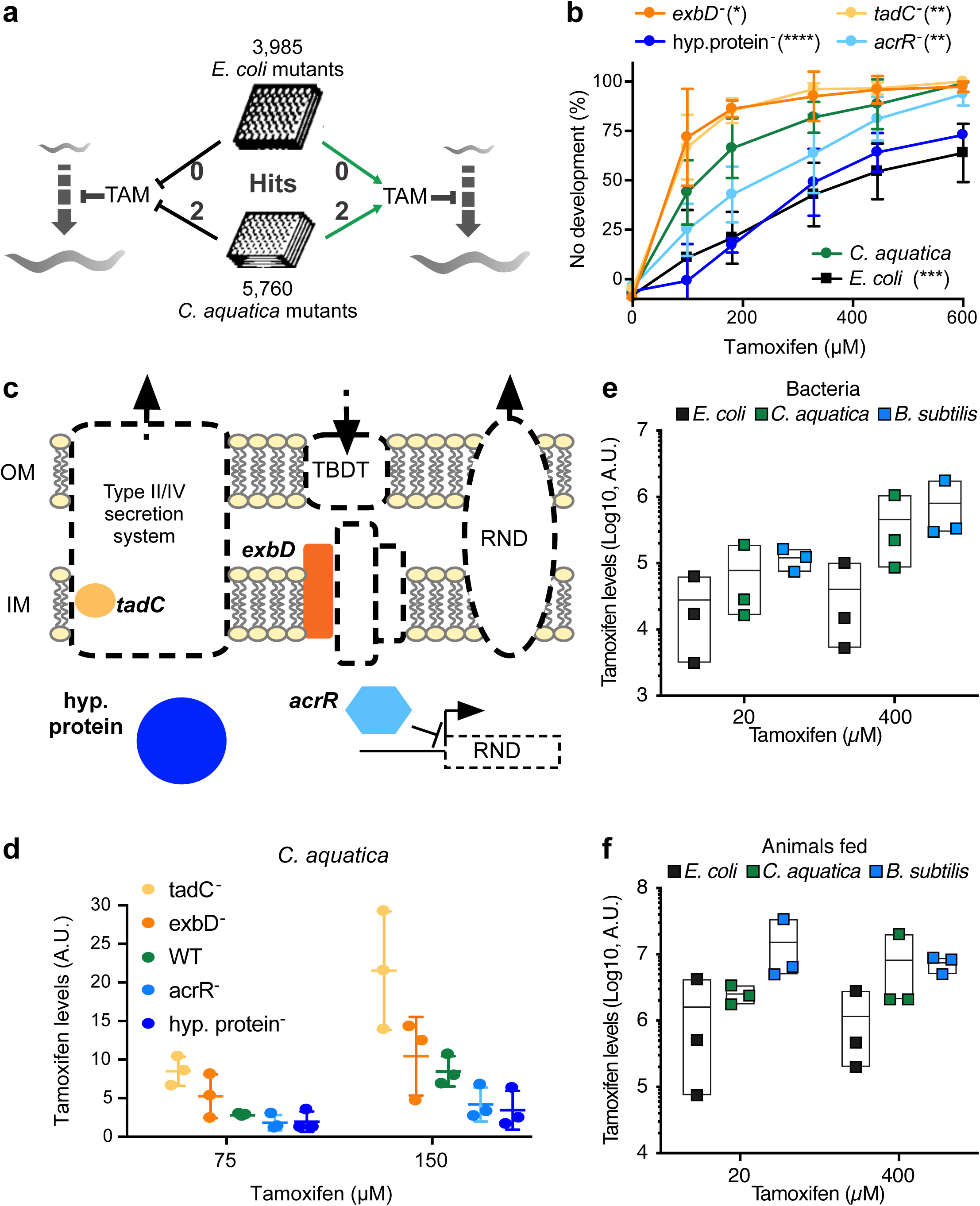
*C. aquatica* transport mutants modulate tamoxifen toxicity in *C. elegans*. **a** Schematic of the bacterial genetics screens performed. **b** DRCs of tamoxifen toxicity in animals fed *C. aquatica* mutants identified in the primary screen **c** Schematic of the function of the genes mutated in the strains identified in **Fig. 3a-b**. Genes encode for factors involved in the type II secretion system (*tadC*), the TonB-dependent transport (*exbD*), or the *acrA/B* multidrug efflux pump (transcriptional repressor *acrR*). **d** Quantification of tamoxifen levels accumulating in *C. aquatica* strains grown on tamoxifen-containing NGM plates. **e** Quantification of tamoxifen levels accumulating in wild-type *E. coli*, *C. aquatica*, and *B. subtilis* strains grown on tamoxifen-containing NGM plates. **f** Quantification of tamoxifen levels in animals fed wild-type *E. coli*, *C. aquatica*, and *B. subtilis* strains in tamoxifen-containing NGM plates.

Next, we compared tamoxifen levels in the three bacterial species, and in animals fed these different diets. We found that bacteria that confer increased toxicity accumulate 5-10-fold more tamoxifen, which translates to an 5-10-fold increased drug accumulation in *C. elegans* (**Fig. 2e**, **f**). However, these differences do not explain the dramatic tamoxifen toxicity in animals fed *B. subtilis*.

Taken together, different bacterial species deliver different amounts of tamoxifen to *C. elegans*, indicating that bioavailability affects, in part, toxicity in the animal. However, these effects are not sufficient to explain the large differences in toxicity depending on bacterial diet, nor do they provide insight into the mechanisms of tamoxifen toxicity in *C. elegans*.

### *C. elegans* fatty acid synthesis protects against tamoxifen toxicity

Because differences in drug bioavailability are not sufficient to explain the differences in tamoxifen toxicity in animals fed the three different bacteria, and since tamoxifen is only toxic to *C. elegans* fed *E. coli* at high micromolar doses (**Fig. 1d**), we hypothesized that *C. elegans* metabolism may affect drug toxicity. To test this, we used RNAi of 1,495 *C. elegans* metabolic genes to identify genes that increased or decreased drug toxicity when knocked down. These include genes with known metabolic functions that make up the genome-scale iCEL1314 metabolic network model, as well as additional genes predicted to encode metabolic enzymes^29^. We did not find any genes that, upon RNAi knockdown, rendered the animals less sensitive to tamoxifen. We did, however, identify three genes for which RNAi knockdown led to increased tamoxifen toxicity (**Fig. 3a**). These genes include *dhs-19*, which encodes a short-chain <u>d</u>e<u>h</u>ydrogena<u>s</u>e of unknown function that localizes to lipid droplets ^30^ and is predicted to function in retinol metabolism ^31^. The other two genes, *elo-3* and *elo-6,* both encode elongases predicted to be involved in long chain FA biosynthesis ^32, 33^. We generated a *dhs-19* deletion mutant by CRISPR/Cas9 genome editing and confirmed that this strain is indeed more sensitive to tamoxifen (**Fig. 3b** and **Supplementary Fig. 3**).

**Fig. 3:**
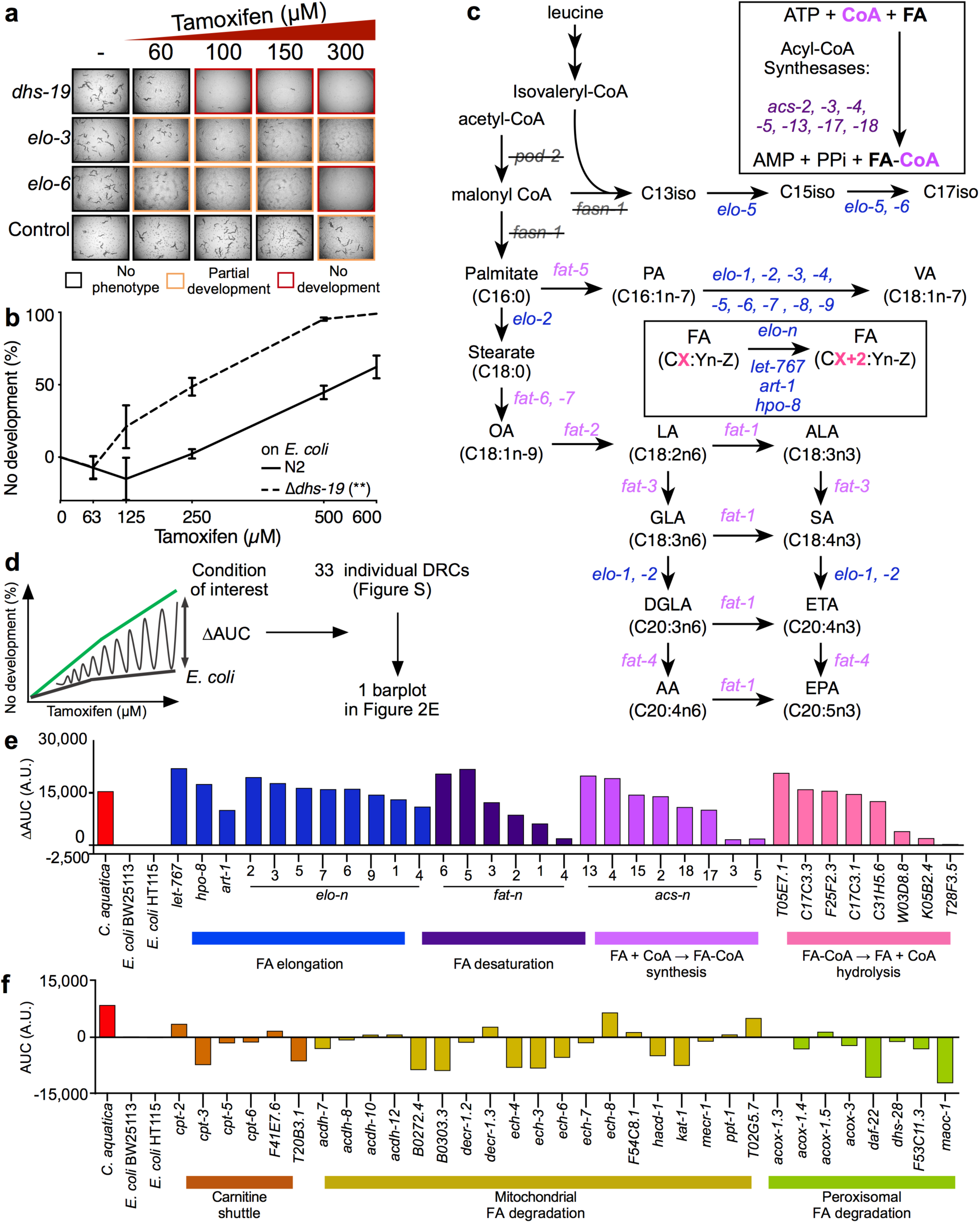
Host long chain fatty acid synthesis modulates tamoxifen toxicity. **a** Screening a metabolic gene RNAi library identified three genes (*dhs-19*, *elo-3* and *elo-5*) that, when knocked down, increase TAM toxicity. Animals were fed *E. coli* HT115 expressing double stranded RNA as indicated. Control indicates *E. coli* containing vector control plasmid (pL4440). Bright-field images were taken at 2x magnification after a 48h exposure to tamoxifen. **b** Tamoxifen dose response curves comparing Δ*dhs-19* mutant animals to wild type animals. Statistical significance was assessed by performing 2way anovas using Graphpad Prism (v9) on original DRCs and is reported in **Supplementary Table 1.** See also **Supplementary Fig. 4**. **c** *C. elegans* fatty acid biosynthesis pathway. Gene names are colored by enzymatic function: FA elongation in blue, FA desaturation in pink, and acyl-CoA synthesis in purple. **d** Cartoon illustrating calculation of differential area under curve (ΔAUC) used in panels E and F. AUCs were calculated using average DRCs. Statistical significance was assessed by performing 2way anovas using Graphpad Prism (v9) on original DRCs and is reported in **Supplementary Table 2**. See also **Supplementary Fig. 5**. **e**, **f** Bar graph showing ΔAUC values obtained from average full DRCs for animals exposed to RNAi of indicated genes.

While mutant strains are available for *elo-3* and *elo-6*, we could not use these animals because their development is strongly impaired even in absence of drug exposure (data not shown). Both *elo-3* and *elo-6* function in FA biosynthesis (**Fig. 3c**). We therefore wondered if these two genes specifically affect tamoxifen toxicity, or if FA metabolism is more broadly involved even though no other genes were captured in the RNAi screen. To test this, we performed full tamoxifen DRC experiments for the knockdown of 33 genes encoding enzymes predicted to be involved in FA elongation and desaturation^33^ (**Fig. 3c**). In addition, we included eight genes encoding acyl-CoA synthetases that catalyze the addition of co-enzyme A (CoA) to FAs and eight genes encoding enzymes that remove CoA. We calculated the difference between the area under the curve (ΔAUC) for each RNAi condition + tamoxifen relative to *E. coli* (**Fig. 3d**). Statistical significance was evaluated on the original DRCs (**Supplementary Fig. 4**, and **Supplementary Tables 1, 2**). Remarkably, knockdown of most FA biosynthesis genes tested significantly increased tamoxifen toxicity (**Fig. 3e**, **Supplementary Fig. 4,** and **Supplementary Table 1**). These results show that *C. elegans* FA synthesis is broadly involved in mitigating tamoxifen toxicity. This points to a systems-level involvement of FA biosynthesis rather than a single FA species.

Next, we asked whether genes involved in mitochondrial and peroxisomal FA degradation also modulate tamoxifen toxicity and found that, of 33 genes tested, only *ech-8* appeared to increase tamoxifen toxicity, but this effect was not statistically significant (**Fig. 3f**, **Supplementary Fig. 5**). Knockdown of 13 of the 33 FA degradation genes tested led to a decrease in tamoxifen toxicity, with only *daf-22* RNAi being statistically significant (**Supplementary Table 2**). This result shows that the biosynthesis, but not degradation, of long chain FAs is critical to mitigate tamoxifen toxicity in *C. elegans*.

### Bacterial diets modulate *C. elegans* fatty acid profiles

Previously, it has been shown that different strains of *E. coli* diets affect the FA composition of *C. elegans*^34^. Since *C. elegans* FA synthesis modulates tamoxifen toxicity, we hypothesized that the three bacterial species used herein may elicit different FA profiles in the animal. To test this, we measured both bacterial and *C. elegans* FA profiles by GC-MS (**Supplementary Tables 3**, **4**). The bacterial FA profiles showed dramatic differences: *E. coli* mainly contained the saturated FAs (SFAs) palmitic acid and stearic acid, cyclopropane FAs (CPFAs), and the mono-unsaturated FAs (MUFAs) oleic acid, palmitoleic acid and cis-vaccenic acid; *C. aquatica* mainly contained the MUFAs oleic acid and palmitoleic acid; and *B. subtilis* almost exclusively contains monomethyl branched chain FAs (mmBCFAs), which agrees with previous findings^35^ (**Fig. 4a**).

**Fig. 4:**
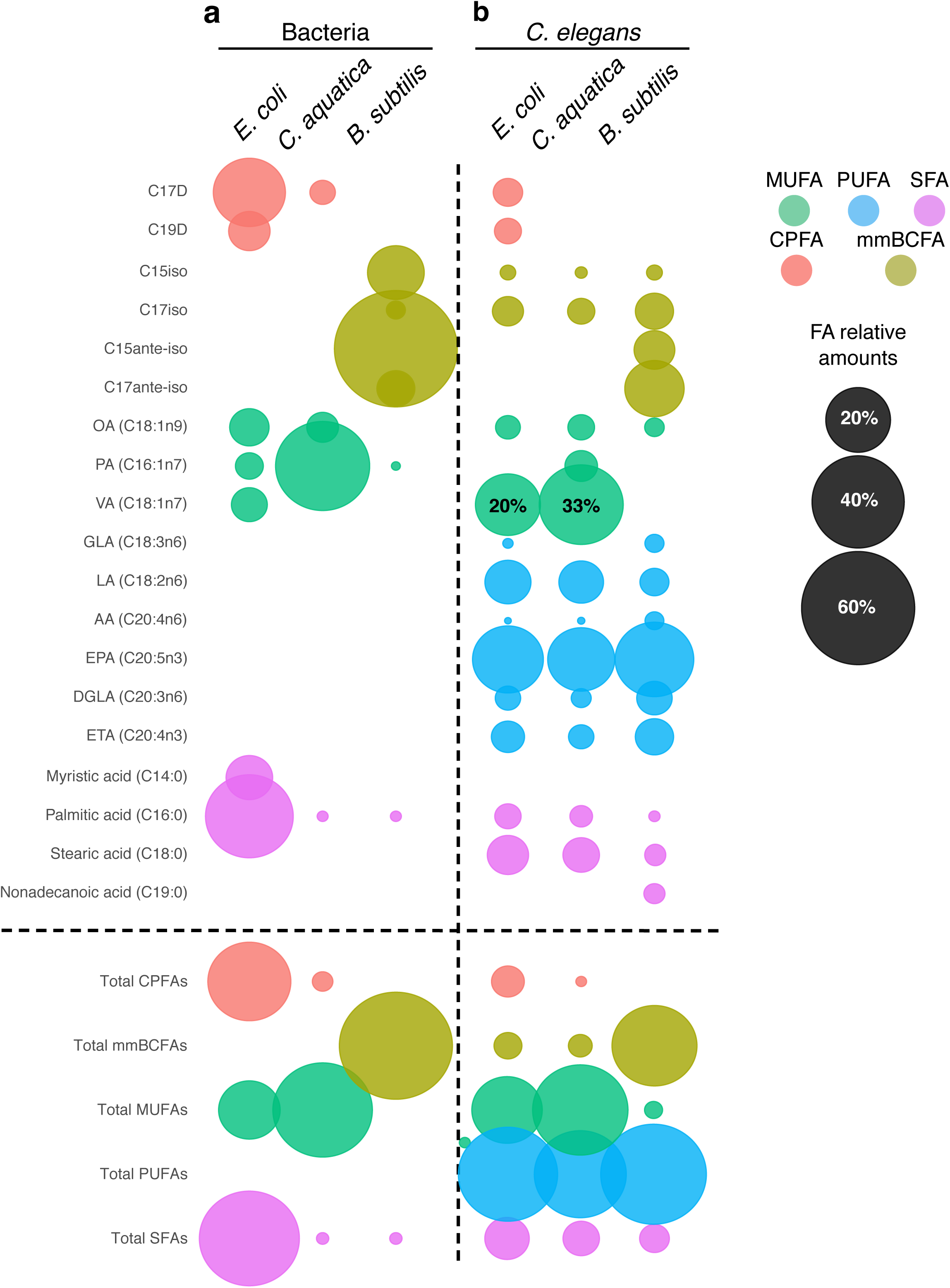
Long chain fatty acid profiles of bacteria and *C. elegans* fed different bacteria. **a, b** Bubble plots showing long chain fatty acid proportions measured by GC-MS in the indicated bacteria (a), and animals fed the indicated bacteria (b). Bubble sizes are indicative of FA ratios within the sample of interest and are not indicative of absolute quantities. Ratios are available in **Supplementary Table 3**, **4**. Data are represented as mean of three to six biological replicates. Statistical pairwise comparisons are available in **Supplementary Table 5**.

The FA profiles of *C. elegans* fed the three bacteria reflected the three bacterial diets in some but not all respects. One striking observation is that the proportion of each of the different poly-unsaturated FAs (PUFAs) is diet-independent (**Fig. 4b**). This indicates that the animal can effectively convert different dietary FAs into PUFAs by elongation and/or desaturation. Oleic acid is the primary substrate for PUFA synthesis and is present in *E. coli* and *C. aquatica* but absent in *B. subtilis* (**Fig. 4a**). This could indicate that PUFAs are generated by a combination of FA metabolic steps that yield this even-chained MUFA from odd-chained mmBCFAs in animals fed *B. subtilis* or synthesizes them de novo.

Another striking observation is that most other types of FAs are quite different in proportion in *C. elegans* fed different bacterial diets. Most notable are the near absence of MUFAs, and presence of C15 and C17-ante-iso mmBCFAs in animals fed *B. subtilis* (**Fig. 4b**). The differences between animals fed the other two bacteria are more subtle, with a greater proportion of the MUFA cis-vaccenic acid in *C. aquatica*-fed animals, relative to those fed *E. coli* (**Fig. 4b**). Altogether, the different bacterial diets result in a different proportion of the different types of FAs: high overall levels of MUFAs in animals fed *E. coli* or *C. aquatica* and high overall levels of mmBCFAs in animals fed *B. subtilis* (**Fig. 4b**, bottom). Taken together, these results show that *C. elegans* maintains its PUFA content in a narrow and well-defined regime, while it can tolerate different proportions of the other types of FAs.

### Dietary fatty acids modulate tamoxifen toxicity

Our findings so far lead us to consider two models: either tamoxifen treatment could alter the animal’s FA profile, leading to diet-dependent toxicity, or differences in *C. elegans* FA composition elicited by bacterial diet may modulate tamoxifen toxicity. To discriminate between these two models, we first measured bacterial and *C. elegans* FAs in the presence or absence of tamoxifen and found that the drug did not modulate the animal’s FA profiles (**Supplementary Table 5**). Thus, we did not find evidence to support the first model.

To evaluate the second model, we first asked whether supplementation of a FA cocktail, containing equimolar amounts of the PUFAs arachidonic and linoleic acid, the SFAs lauric and myristic acid, and the MUFA oleic acid^36^, would modulate tamoxifen toxicity in animals fed either of the three bacterial diets. We found that FA cocktail supplementation suppressed tamoxifen toxicity but only in animals fed *C. aquatica*; it had no effect on tamoxifen toxicity in animals fed *E. coli* or *B. subtilis* (**Fig. 5a-c**). Next, we tested the effects of the supplementation of individual FAs and found that MUFAs such as cis-vaccenic acid increased tamoxifen toxicity on animals fed *E. coli*, but not *C. aquatica* (**Fig. 5d, e**, and **Supplementary Fig. 6**). Since *C. aquatica*-fed animals contain more cis-vaccenic acid than *E. coli*-fed animals (33% vs 20%, **Fig. 4b**), this suggests that this difference may contribute to increased tamoxifen toxicity on the *C. aquatica* diet.

**Fig. 5:**
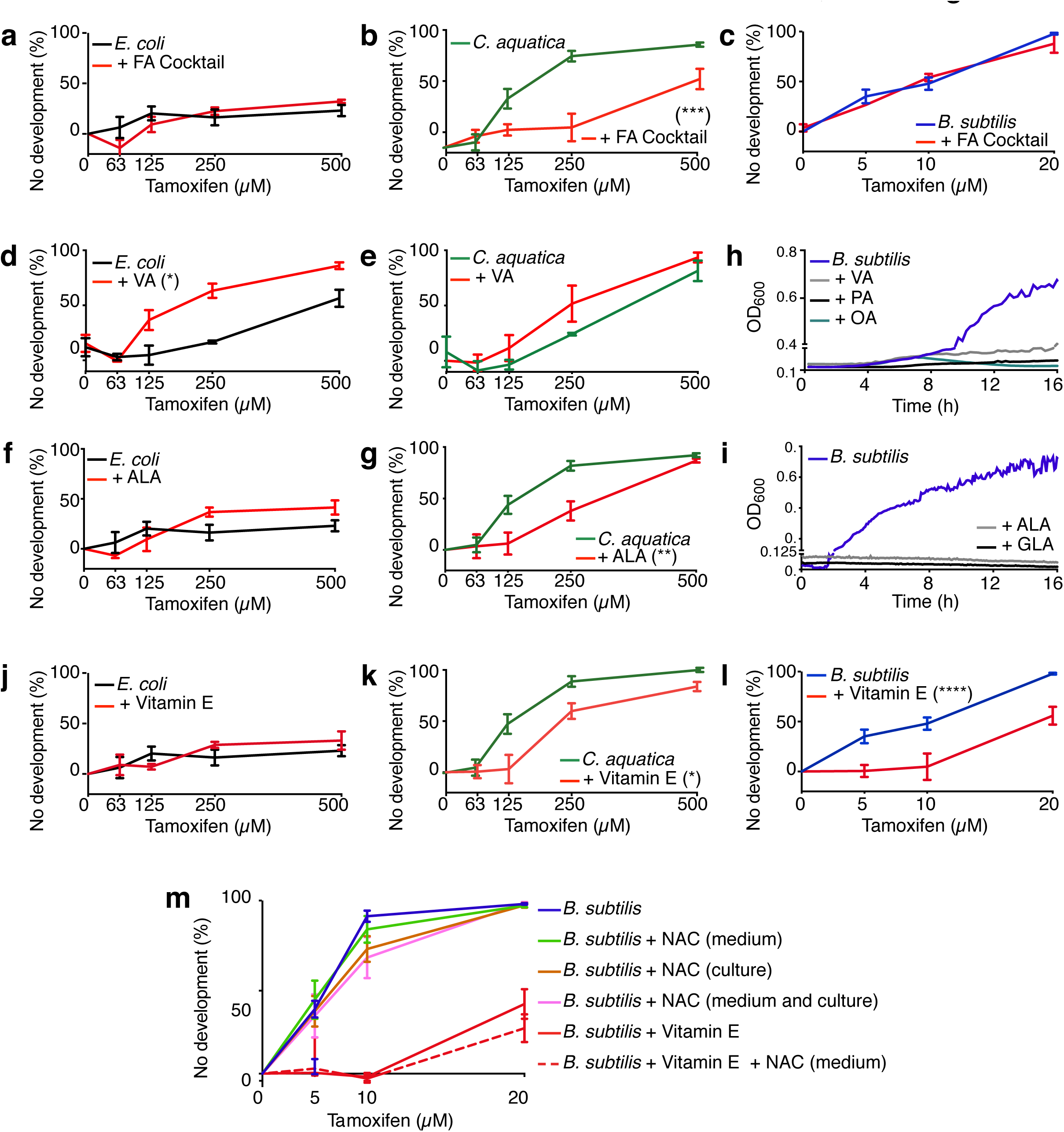
Long chain fatty acid supplementation modulates tamoxifen toxicity. **a**-**m** Unless otherwise indicated, fatty acids and vitamin E were supplemented to bacterial culture medium, as described previously (Devkota et al., 2021). **a**-**c** DRCs of tamoxifen toxicity with and without FA cocktail supplementation on animals fed *E. coli* (**a**), *C. aquatica* (**b**) or *B. subtilis* (**c**). **d**, **e** DRCs of of tamoxifen toxicity with and without PUFA (ALA) supplementation on animals fed *E. coli* (**d**) or *C. aquatica* (**e**). **f** Growth curves of *B. subtilis* supplemented with PUFAs ALA or GLA. For clarity, no error bars are reported in the graph, n=3. **g**, **h** DRCs of tamoxifen toxicity with and without MUFA (VA) supplementation on animals fed *E. coli* (**g**) or *C. aquatica* (**h**). **i** Growth curves of *B. subtilis* supplemented with MUFAs VA, PA, or OA. No error bars are reported in the graph, n=3. **j**-**l** DRCs of tamoxifen toxicity with and without vitamin E supplementation on animals fed *E. coli* (**j**), *C. aquatica* (**k**) or *B. subtilis* (**l**). **m** DRCs of tamoxifen toxicity with and without the antioxidants vitamin E and N-acetyl-cysteine (NAC). Culture: NAC was supplemented in the bacterial culture medium as for vitamin E. Medium: NAC was supplemented to the NGM. DRC: dose response curve, FA: fatty acid, MUFA: mono-unsaturated FA, PUFA: poly-unsaturated FA, ALA: alpha-linoleic acid, GLA: gamma-linoleic acid, DGLA: di- homo-gamma-linoleic acid, VA: cis-vaccenic acid, PA: palmitoleic acid, OA: oleic acid. Statistical analysis of DRCs was conducted by performing 2-way Anova using GraphPad Prism (v9) and are reported in **Supplementary Table 6**. Larger panel of fatty acid supplements was tested in **Supplementary Fig. 7.**

Supplementation of individual PUFAs such as α-linoleic acid decreased tamoxifen toxicity on animals fed *C. aquatica* (**Fig. 5f, g**, and **Supplementary Fig. 6**). On an *E. coli* diet PUFAs had no effect. However, animals are generally tolerant to tamoxifen, which limits the ability to observe decreased tamoxifen toxicity. Since the proportion of PUFAs is similar in animals fed each of the three bacterial diets (**Fig. 4b**), this indicates that the ratio of PUFAs to other FAs is more relevant than absolute levels. In contrast to *E. coli* and *C. aquatica*, supplementation of either MUFAs or PUFAs abolished the growth of *B. subtilis*, and therefore these FA could not be tested for tamoxifen toxicity modulation in *C. elegans* fed these bacteria (**Fig. 5h, i** and **Supplementary Fig. 7**). Together with the observation that knockdown of genes involved in FA synthesis increases tamoxifen toxicity, these results point to a complex interplay between dietary FA, FA synthesis and tamoxifen toxicity.

We next asked whether bacterial metabolism was required for the effects of supplemented FAs on tamoxifen toxicity. We generated metabolically inactive bacterial powders by mechanical lysis followed by lyophilization as previously described^7^, and seeded these powders onto tamoxifen-containing plates. We observed that the dietary-dependent effects were maintained: *E. coli* powder-fed animals developed in presence of 600 µM tamoxifen, while animals fed *C. aquatica* powder did not. Similar to feeding live *C. aquatica*, supplementation of FA cocktail allowed animals to develop in presence of 600 µM tamoxifen when fed metabolically inactive *C. aquatica* (**Supplementary Fig. 8**). *B. subtilis* powder did not support *C. elegans* development and therefore could not be tested for tamoxifen toxicity (**Supplementary Fig. 8**). These results indicate that active bacterial metabolism does not affect tamoxifen toxicity in *C. elegans*, which is also supported by the lack of metabolic gene mutants identified in the bacterial mutant screens. Instead, these observations further support the idea that different bacteria provide the animal with different FAs and that these FAs differently modulate tamoxifen toxicity.

*E. coli*-fed animals harbor high levels of CPFAs, while *B. subtilis*-fed animals contain high levels of mmBCFAs (**Fig. 4b**). Therefore, we asked whether these FAs could modulate tamoxifen toxicity in *C. elegans*. We used bacterial genetics to assess the importance of CPFAs because they are exclusively synthesized in bacteria by CFA enzymes^37^. Because CPFAs specifically accumulate in *E. coli* (**Fig. 4b**), and because this diet renders the animals less sensitive to tamoxifen toxicity, we reasoned that CPFAs may be protective. We fed *C. elegans* two independent *E. coli* Δ*cfa* strains from the Keio library ^24^ and did not observe and effect on tamoxifen toxicity (**Supplementary Fig. 9**). Next, we tested whether mmBCFAs, which specifically accumulate in *B. subtilis*, would enhance tamoxifen toxicity. mmBCFAs occur in two types, iso-mmBCFAs and ante-iso-mmBCFAs^38^. We were not able to test mmBCFAs by supplementation due to cost and supply issues. However, because ante-iso mmBCFAs are synthesized from isoleucine supplied in growing media, we could assess their contribution to tamoxifen toxicity by growing *B. subtilis* in the absence of isoleucine^38^. There was no difference in tamoxifen toxicity in animals fed *B. subtilis* grown in LB (rich in isoleucine) or minimal M9 media supplemented with glucose as carbon source (**Supplementary Fig. 10**). While not unequivocally ruling out the contribution of mmBCFAs, these data show that a switch from ante-iso to iso-mmBCFAs in the food uptake does not modulate tamoxifen toxicity.

Previously, it has been shown that tamoxifen inhibits glucosylceramide synthesis in human cells^39^. In *C. elegans*, mmBCFAs are converted into the sphingolipid d17- isoglucosylceramide by ELO-5, and this conversion is important to support development and growth ^40^. Since we found that *elo-5* RNAi enhances tamoxifen toxicity (**Fig. 3e**), we next asked whether tamoxifen elicits a d17-isoglucosylceramide deficiency that explains drug toxicity. In *C. elegans*, knockdown of *elo-5* can be rescued by combining it with a mutation in components of the NPRL2/3 complex^40^ (**Supplementary Figure 11**). If tamoxifen toxicity is due to the lack of d17-isoglucosylceramide, we would expect that mutants in the NPRL-2/3 complex would mitigate toxicity. However, we found that *nprl-3* deletion mutants showed equal sensitivity to tamoxifen as wild type animals (**Supplementary Fig. 11**). This result shows that tamoxifen does not affect *C. elegans* by depletion of d17-isoglucosylceramide.

### How do FAs and FA biosynthesis modulate tamoxifen toxicity in *C. elegans*?

Since FAs are easily oxidized and rapidly turnover in cellular and organellar membranes^41^, we hypothesized that tamoxifen treatment causes FA oxidation, and that this oxidation is mitigated by replenishing oxidized FAs with newly synthesized FAs. Vitamin E is a potent antioxidant that inhibits FA oxidation and that stabilizes the FA cocktail^42^. We found that vitamin E had no effect on tamoxifen toxicity in animals fed either *E. coli* or *C. aquatica*, indicating that it is the FAs in the FA cocktail that mitigate toxicity in animals fed *C. aquatica*. However, vitamin E supplementation did suppress tamoxifen toxicity in animals fed *B. subtilis* (**Fig. 5j-l**), when N-acetylcysteine (NAC), another antioxidant that can rescue the growth of *C. elegans* on iron-deficient *E. coli* mutants^43^ did not affect tamoxifen toxicity (**Fig. 5m**). These results indicate that FA oxidation, but not the general accumulation of reactive oxygen species (ROS), affects tamoxifen toxicity in *C. elegans* when fed *B. subtilis*. Taken together, these results indicate that FA or vitamin E supplementation can mitigate tamoxifen toxicity differently depending on which bacteria the animals are fed, which suggests that different, diet-dependent toxicity mechanisms are involved.

### Tamoxifen induces the expression of type II detoxification genes

We next used expression profiling by RNA-seq to ask whether tamoxifen supplementation modulates the expression of genes involved in FA metabolism or oxidative stress in *C. elegans*. To avoid capturing mRNA changes due to differences developmental rate, we used a tamoxifen dose at which no developmental arrest was observed, but in which animals are delayed by ∼12 hours. The development of animals exposed to tamoxifen was monitored hourly, and samples were collected 12h after control animals. A total of 558 and 738 genes were up- and down regulated, respectively (>2-fold change, P< 0.01, **Supplementary Table 6**). Tamoxifen supplementation did not change the expression of genes involved in FA biosynthesis or degradation genes with the exceptions of *fat-5,* which increased by three-fold, and *ech-6* which was reduced by ∼two-fold (**Fig. 6a**). Together with the observation that tamoxifen supplementation did not alter the animal’s FA profiles, these results show that tamoxifen does not modulate FA metabolism. Instead, it suggests that the animal’s FA state modulates tamoxifen toxicity.

**Fig. 6:**
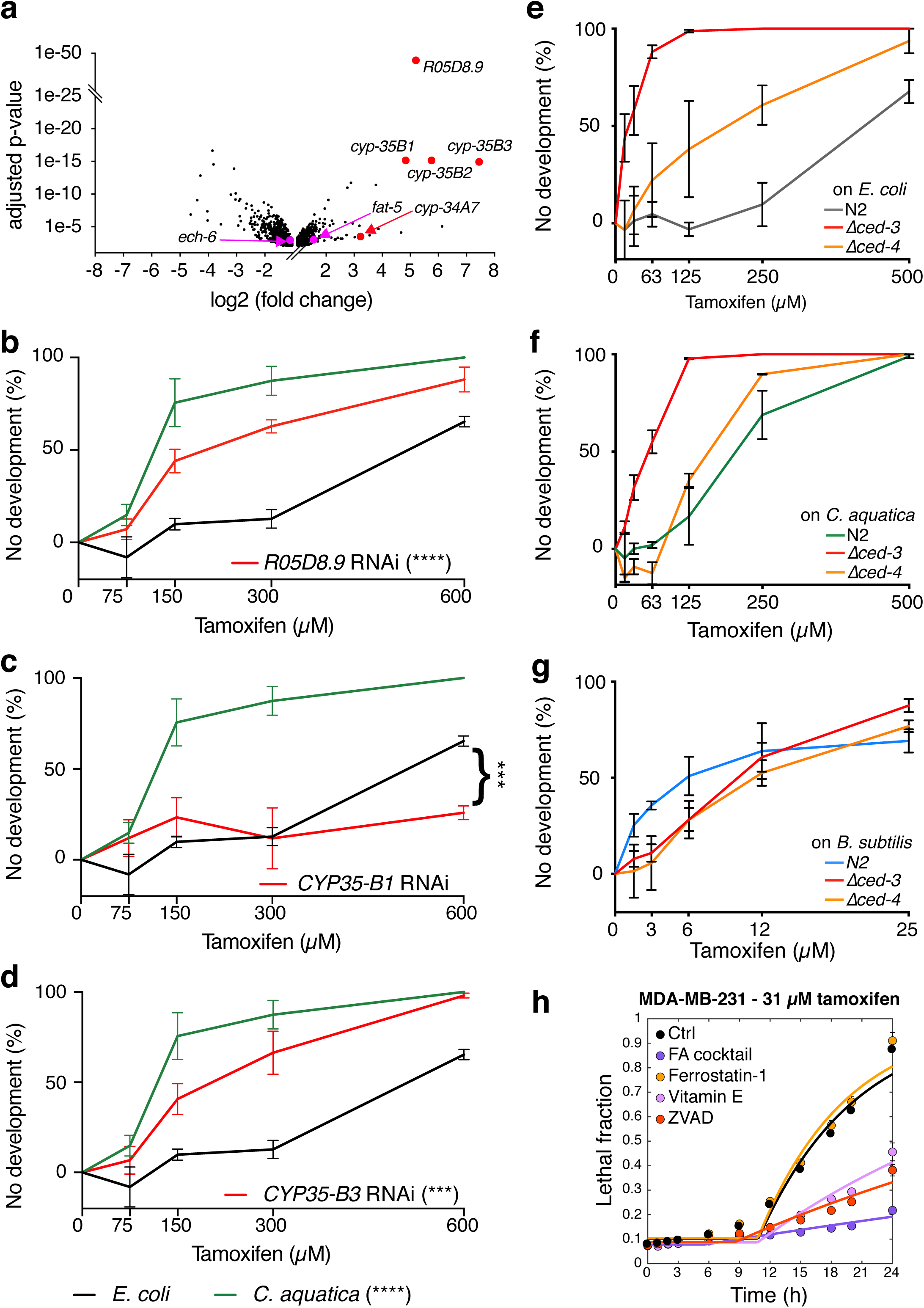
Bacteria affect the modulation of tamoxifen toxicity by apoptosis. **a** Volcano plot distribution of genes which expression changed significantly in animals fed *E. coli* and exposed to 400 µM tamoxifen (FDR<0.1, >2-fold change). Full list is provided in **Supplementary Table 6**. Pink indicates genes encoding enzymes participating in FA metabolism, and red indicates genes encoding for enzymes that may be involved in the detoxification of tamoxifen. See also **Supplementary Fig. 12**. **b-d** DRCs of tamoxifen toxicity in animals fed *E. coli* expressing double stranded RNA as indicated. Control indicates *E. coli* containing vector control plasmid (pL4440). Error bars represent SEM, n=3. Statistical analysis of DRCs was conducted by performing 2-way Anova using GraphPad Prism (v9). Notably, the significance of *CYP35-B1* RNAi (**c**) rescue was not captured on the entire curve where no substantial toxicity can be overserved on the *E. coli* control, but rather on the highest drug-concentration where reliable toxicity can observed. **e-g** DRCs of tamoxifen toxicity on apoptosis deficient mutant animals fed *E. coli* (**e**), *C. aquatica* (**f**) or *B. subtilis* (**g**). **h** Kinetics of tamoxifen toxicity in ER-negative (MDA-MB-231) breast cancer cells in presence of absence of FA cocktail, Ferrostatin-1 (ferroptosis inhibitor), Vitamin E, or ZVAD (apoptosis inhibitor). See also **Supplementary Fig. 13**.

**Fig. 7:**
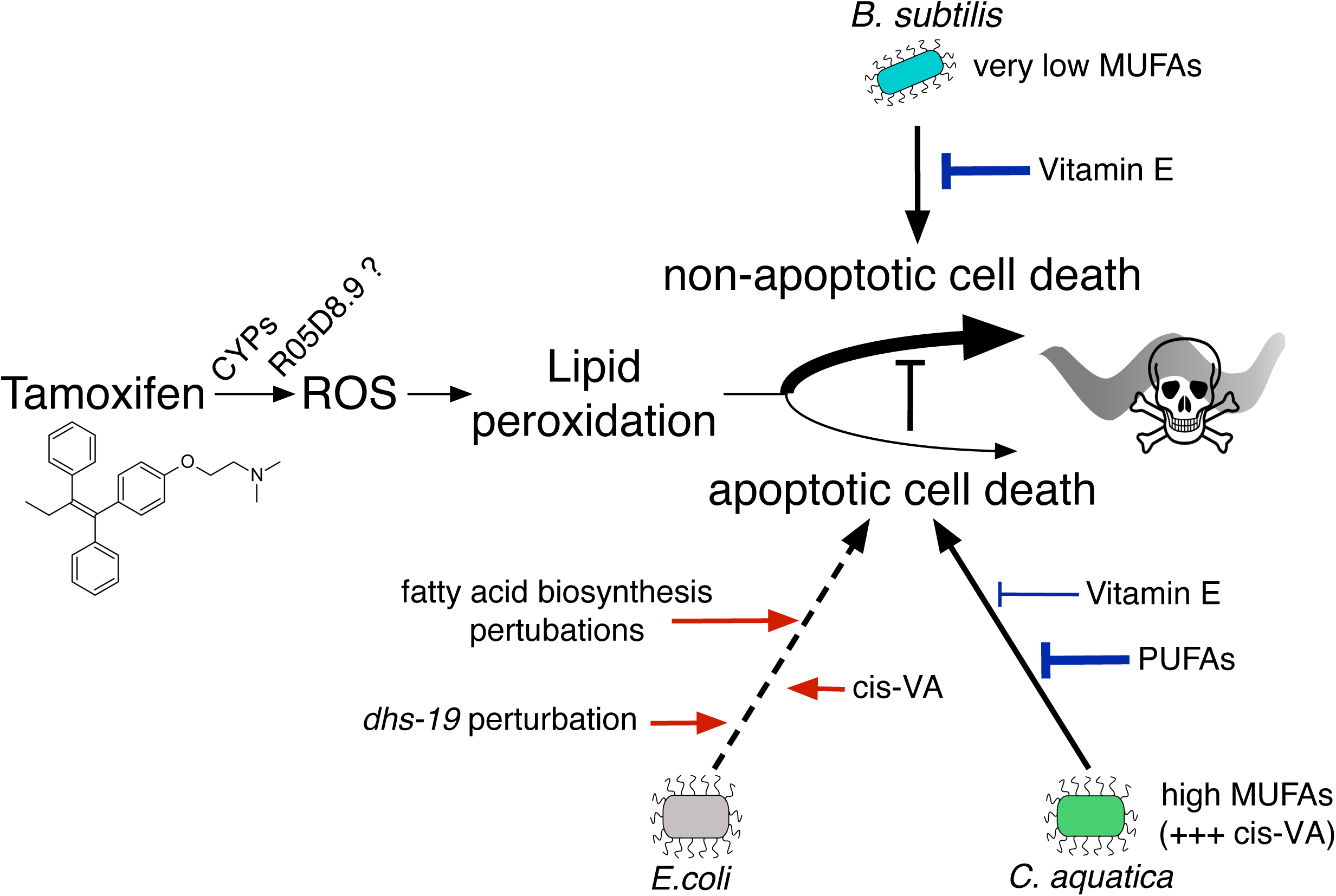
Model of Interplay Between Bacteria, Fatty Acid Metabolism, and Death Pathways Affecting Tamoxifen Toxicity.

Several genes that change in expression in response to tamoxifen supplementation pointed to potential mechanisms of toxicity. First, tamoxifen induced the expression of R05D8.9 (38-fold, adjusted *p-*value: 1.2x10^-49^), which encodes a predicted 17beta-hydroxysteroid dehydrogenase 14 (17βHSD14) homolog (**Fig. 6a** and **Supplementary Table 6**). In humans, 17βHSD14 enzymes are associated with the response to tamoxifen in breast cancer^44^. These enzymes are further known to interconvert different steroids. We tested whether the induction of this gene is functional and found that R05D8.9 knockdown increased tamoxifen toxicity (**Fig. 6b**). This observation suggests that R05D8.9 is involved in the detoxification of tamoxifen. The expression of several CYP genes (*cyp-35B3, -35B2, -35B1,* and *-34A7*), which encode cytochrome P450 enzymes involved in type II detoxification, is induced by tamoxifen supplementation (179-, 55-, 29-, and 10-fold, adjusted *p-*values: 1.6x10^-15^, 9.2x10^-16^, 9.2x10^-16^ and 4.1x10^-4^) (**Fig. 6a** and **Supplementary Table 6**). Cytochrome P450 enzymes have been reported to modify tamoxifen in humans^45^, and are a source of oxidative stress as their enzymatic cycle involves significant leakage of electrons ^46^. Again, we tested whether these expression changes are functional and found that knockdown of *cyp-35B3* increased drug tamoxifen toxicity, while knockdown of *cyp-35B1* decreased toxicity at high doses (**Fig. 6c, d**). These results demonstrate the functional involvement of CYP-450 enzymes in detoxifying tamoxifen toxicity, and further suggest, that different CYP-450 proteins can exhibit differential tamoxifen- detoxifying and ROS-generating potentials, which is in agreement with previous observations^45, 47^. Taken together, tamoxifen does not transcriptionally rewire FA metabolism. Instead, it induces the expression of ROS-generating detoxifying enzymes, which may contribute to FA oxidation.

### Bacteria elicit different tamoxifen-induced death mechanisms

Lipid metabolism has been linked to different mechanisms of cell death, especially apoptosis and ferroptosis^48, 49^. In fact, it has been shown that cell survival is dependent on lipid homeostasis, and both excessive MUFAs or SFAs have been shown to be pro-apoptotic, depending on the cell lines^49^. Further, several MUFAs have been reported to protect against ferroptosis^50^, which is characterized by lipid peroxidation, while others, specifically cis-vaccenic acid have been reported to induce cell death^51^. PUFAs, most notably di-homo-gamma-linoleic acid, have been shown to induce ferroptosis in both human cells and in *C. elegans*^52^. Given these connections, we asked whether apoptosis contributes to tamoxifen toxicity in *C. elegans*. To test this, we examined tamoxifen toxicity in *ced-3* and *ced-4* mutants that are deficient in apoptosis^53^. CED-3 is a caspase that is essential for apoptosis, and CED-4 is an activator of CED-3. Remarkably, loss of *ced-3*, and to a lesser extent *ced-4,* greatly increased sensitivity to tamoxifen in animals fed either *E. coli* or *C. aquatica* but had no effect on animals fed *B. subtilis* (**Fig. 6e-g** and **Supplementary Fig. 12**). This result suggests that, when fed *E. coli* or *C. aquatica,* tamoxifen toxicity employs apoptosis, and that *ced-3* and *ced-4* mutant animals switch to another, more potent mode of death that is suppressed when apoptosis is functional. We found that a high dose of vitamin E could mitigate tamoxifen toxicity in *ced-3* mutant animals fed *C. aquatica* but not *E. coli* (**Supplementary Fig. 12**). This indicates that, in the absence of *ced-3*, tamoxifen toxicity in animals fed *C. aquatica* involves FA oxidation, similar to what we observed with wild type animals fed *B. subtilis*.

Finally, we tested the effects of Vitamin E and FA cocktail on a dose of tamoxifen that is toxic independently of ER in human cells (31 µM, **Fig. 1a, b**). Tamoxifen toxicity was severely suppressed by either supplement in both ER-negative MDA-MB-231 and ER-positive cells (**Fig. 6h**, **Supplementary Fig. 13**). Furthermore, ZVAD, a pan-caspase inhibitor that prevents apoptosis, reproducibly lowered the drug toxicity in MDA-MB-231 but not in T-47D cells. This suggests that, like in *C. elegans*, the ER-independent toxicity of tamoxifen can be elicited through different cell death pathways in human cells.

## Discussion

In this study, we present *C. elegans* as a model to study off-target, ER-independent, mechanisms of tamoxifen toxicity using concentrations in the range of drug-levels accumulating in patients^54, 55^. We found that FA biosynthesis and composition of the animals are major drivers of tamoxifen toxicity, and that this toxicity relies on the potential of the drug to induce oxidative stress and lipid peroxidation. Moreover, we found that both the mechanism and the degree of tamoxifen toxicity are greatly modulated by bacteria, where *B. subtilis* induces a non-apoptotic mechanism that is three orders of magnitude more severe than apoptotic mechanism elicited by tamoxifen in the presence of either *E. coli* or *C. aquatica* (**Fig. 6**).

In the *C. elegans* model, bacteria form a source of nutrition, can be pathogenic, and can inhabit the gut as the animal ages^56^. Both in the laboratory and in the wild, the animals are constantly exposed to bacterial metabolism which provides nutrients and contributes to animal physiology^57^. Our findings illuminate the possible effects the microbiota may have on tamoxifen off-target effect toxicity and the treatment of ER- independent malignancies^17^. It is tempting to speculate that tamoxifen activity could be modulated by perturbations of the microbiota using antibiotics or probiotics. Notably, *B. subtilis*, which is present in a large proportion of commercially available probiotics^58^ may affect patients treated with tamoxifen. Our findings of the role of FA metabolism and oxidative stress further suggest that studies of nutritional intake of fats and vitamin E in humans may shed light on whether the off-target mechanisms discovered herein are conserved.

We uncovered complex relationships between bacteria, host FA metabolism, apoptosis, and tamoxifen toxicity. The literature about the relationship between FAs and cell survival integrates two major components: distinct FA-species display different oxidation potentials^59^, and different potentials to induce alternative cell-death mechanisms^49, 50^. Our data using lipophilic ROS-scavenging vitamin E and apoptosis-deficient animals suggest that tamoxifen-toxicity relies on these two entangled components. MUFAs are generally described as ‘healthy’ FAs, as they buffer the oxidative stress of ROS-sensitive PUFAs^50^. To our knowledge, there is only one report on the effects of the MUFA cis-vaccenic acid, which found that its accumulation induces cell death, when other MUFAs tested in parallel do not^51^. Overall, our observation of extreme tamoxifen toxicity in *B. subtilis-*fed animals that do not provide the animal with MUFAs, and in *C. aquatica-*fed animals that accumulate cis-vaccenic acid are consistent with the literature. Further studies are required to systematically determine the effects of different FAs and lipids on cell survival and tamoxifen toxicity in human cells.

Knockdown of or mutations in *dhs-19* enhance tamoxifen toxicity in animals fed *E. coli*. The precise molecular or metabolic function of DHS-19 is unknown. It encodes a dehydrogenase and is a marker of lipid droplets^60^, which are essential for FA homeostasis in all eukaryotes^61^. Based on enzyme homology, DHS-19 is predicted to act in retinoid metabolism, through the interconversion of retinol and retinal^33^. This reaction precedes the synthesis of retinyl palmitate which consumes palmitoyl-CoA, which is a precursor for the synthesis of palmitoleic and vaccenic acid, both of which potentiate tamoxifen toxicity in *C. elegans* when animals are fed *E. coli*. Retinoids are also potent ROS scavengers and have been shown to act synergically with vitamin E to reduce lipid oxidative stress^62^. Together, this suggests that the increase in tamoxifen toxicity by *dhs-19* perturbation could result from a complex integration of metabolic cues involving oxidative stress and retinoid metabolism.

*C. elegans* is a remarkable animal with a fixed lineage in which wild type animals develop deterministically to adults with precisely 959 somatic nuclei^63, 64^. Therefore, one would assume that all adult animals are essentially the same. Here we show that wild type animals can exhibit marked differences in their FA composition, depending on which bacterial diet they are fed. To our knowledge, *C. elegans* FA composition has only been studied in animals fed *E. coli*, and the profiles observed in our experiments are consistent with the only available previous study^34^. Here, we observe much more drastic changes in FA composition in animals fed *C. aquatica* or *B. subtilis*, relative to those fed *E. coli*. First, the MUFA, mmBCFA or CPFAs content of the animals strongly reflects the dietary input; a diet of *C. aquatica* is rich in MUFAs, and a diet of *B. subtilis* is rich in mmBCFAs, and these FAs accumulate in the animals. This result demonstrates that the animal’s membranes are permissive to varying levels of these types of FAs. In contrast, the PUFA composition of *C. elegans* is largely independent of its bacterial diet. While PUFAs are essential dietary FAs in humans, *C. elegans* synthesize PUFAs de novo^65^. This result indicates that *C. elegans* PUFA levels are sensed, and that metabolic fluxes in FA biosynthesis pathways are adjusted to maintain relative levels in a tight regime. It is therefore perhaps not surprising that PUFA deficiencies have been associated with developmental and neuronal defects in *C. elegans*^66^. Future studies are required to reveal to what the extent wild type *C. elegans* exhibit other fitness changes in response to differences in FA composition.

How does cell-death connect to organism death? Fifty years after Horvitz and Sulston first described apoptosis in *C. elegans*^63^, it remains unclear how cell death mechanisms communicate and how they relate to organismal phenotypes including death. Other open questions that remain to be answered are: What is the exact nature of the triggers directing decisions in the induction of alternative cell-death mechanisms? And how do bacteria affect cell-death potentials? This study provides a foundation for future studies to address fundamental questions related to cell-death and organismal fate.

## Methods

### C. elegans

*C. elegans* N2 (Bristol) was used as the wild-type strain. Prior to all experiments, animals were maintained on Nematode Growth Media (NGM) and fed a diet of *E. coli* OP50. To prepare L1-synchronized animals, eggs purified by dissolving gravid animals in NaOH-buffered bleach were collected and incubated for 20h in M9 buffer for hatching. The *C. elegans ced-3(n717)* mutant strain was kindly provided by the Francis lab (UMass Medical School) and was first described in (8598288). The *C. elegans dhs- 19(VL1313)* mutant strain was generated using CRISPR-Cas9 genome editing^67^.

### Bacterial strains

*E. coli* BW25113, *E. coli* HT115, *C. aquatica* DA1877, and *B. subtilis* sp168 were grown in LB, overnight at 37°C to stationary phase under agitation (200 rpm). Alternatively, *B. subtilis* sp168 was grown in M9 at 37°C to stationary phase (∼24h). The dietary effects on tamoxifen toxicity described herein were independent of the *E. coli* strain tested. Experiments were conducted using *E. coli* BW25113, unless stated otherwise. Bacterial powders were generated by mechanical lysis followed by desiccation as previously described^7^. Powders were seeded onto tamoxifen containing NGM plates supplemented antibiotics and prepared without peptone.

### Bacterial genetic screens

Bacterial genetic screens were performed in 96-well plates containing NGM supplemented with 200 and 300 µM tamoxifen for *E. coli* mutants, 100 and 300 µM tamoxifen for *C. aquatica* mutants. Wells were seeded with individual mutant strains, and approximately twenty L1-arrested animals were added to each well on the following day. After 2 and 3 days at 20°C, hits were scored visually using a dissecting microscope as bacterial mutants leading to a difference in animal drug response relative to wildtype-fed animals. Primary hits were then retested in 48-well plates, in two independent experiments, on NGM supplemented with 0.5% DMSO, 50, 100, 150, 300, or 600 µM tamoxifen for *E. coli* and *C. aquatica* mutants. Bacterial mutants that showed robust differences on host drug response were genotyped. *E. coli* and *C. aquatica* and mutants were genotyped using gene-specific primers flanking -200bp and +200bp relative to the start codon and semi-random two step PCR, respectively.

### Tamoxifen-containing NGM plate preparation

Tamoxifen citrate (Selleckchem, S1972) was dissolved to 150 mM in sterile dimethyl sulfoxide (DMSO), aliquoted and stored at -20°C. Stock solutions were diluted in series to 200X the final concentration and added at a final volume of 0.5% to NGM agar heated to 55°C. Drug containing NGM plates were dried overnight at room temperature. NGM was supplemented with antibiotics for experiments in 96-well plate format, and 2 mM IPTG for RNAi experiments. 96-well and 48-well plates were seeded with 15 µL and 30 µL of bacterial cultures, respectively. Plates were then dried for 6h at room temperature and kept overnight at room temperature before being used. *E. coli* cultures were concentrated 1-2X, and *C. aquatica* DA1877 and *B. subtilis* sp168 strains were concentrated 2-4X prior to seeding onto NGM agar plates, to normalize for post-seeding bacterial growth.

Importantly, batch-to-batch effects were observed as the study was conducted with differences in drug toxicity depending on the batch. However, dietary differences on tamoxifen drug toxicity were systematically observed.

### Tamoxifen measurements

Tamoxifen levels were measured on an Agilent 7890B/5977B single quadrupole GC-MS equipped with an HP-5ms Ultra Inert capillary column. Briefly, 5,000 animals were mechanically lysed in 80% methanol using a FastPrep-24 bead beater (MP Biomedicals). Lysates were centrifuged for 10 min at 20,000 × g, and 250 μL of the supernatants were dried using a SpeedVac concentrator SPD111V (Thermo Fisher Scientific). Derivatization was performed by incubating the extracts with methoxyamine hydrochloride (Sigma-Aldrich) [37°C - 1.5h], and then with N-methyl-N- (trimethylsilyl)trifluoroacetamide (Sigma-Aldrich) [37°C, 3h and then 25°C, 5h]. Peak areas were quantified using samples within a linear response range, after mean normalization to total metabolites and blank subtraction. Sample size biases were then corrected by normalizing tamoxifen signals to the metabolomic weight of the sample they originate from (i.e., the sum of all 23-metabolites measured in the run of interest).

For bacteria, lawns were recovered 24h post-seeding, which corresponds to the time at which animals are exposed to tamoxifen in the experiments presented in this manuscript. For *C. elegans*, animals were first grown to the young-adult stage in absence of tamoxifen, on *E. coli*, *C. aquatica* or *B. subtilis*. They were then transferred onto tamoxifen containing plates, seeded with the same bacteria 24h before. Animals were recovered after 12h of exposure.

### RNAi screen

We tested a collection of 1,645 individual *E. coli* HT115 strains expressing dsRNA against a unique *C. elegans* metabolic gene^29^. The screens were performed using 96- well plates containing NGM supplemented with tamoxifen, 50 µg/mL ampicillin, and 2 mM IPTG. We used a low dose of 300 µM tamoxifen to identify knockdowns that aggravate tamoxifen toxicity, and a high dose of 600 µM tamoxifen identify knockdowns that mitigate tamoxifen toxicity. Approximatively 20 L1-arrested animals were added to each well. Two days later, hits were scored visually using a dissecting microscope as knockdowns leading to a difference in animal drug response relative to animals fed an *E. coli* strain containing the empty dsRNA vector backbone (pL4440).

#### *C. elegans* dose-response curves

Dose-response curves (DRCs) were performed in 48-well plates with six to twelve technical replicates. Fifty to one hundred L1-arrested animals were plated onto each well and incubated at 20°C for 48 hours. Animals that developed beyond the L1 stage were counted under a dissecting microscope. The percentage of animals that did not developed was calculated relative to the total number of animals developed in the control condition. For ΔAUCs in **Fig. 3d-f**, we used the average AUCs from the DRCs of the *E. coli* BW25113 and *E. coli* HT115 pL4440 control conditions.

### Bacterial growth curves

Bacteria were first grown overnight in LB at 37°C and shaking at 200 rpm. Next, bacterial cultures were diluted 1:200 and grown in LB supplemented with 1% of FA supplements (or ethanol vehicle as a control). Cultures were then grown for 20h at 37°C and shaking at 200 rpm, in an Eon™ Microplate Spectrophotometer (BioTek) and absorbance at 600 nm was measured every 900 s.

### Fatty acid and vitamin E supplementations

The FA cocktail contains equimolar amounts of the PUFAs arachidonic and linoleic acid, the SFAs lauric and myristic acid, and the MUFA oleic acid^36^ (Sigma-Aldrich, F7050). Undiluted fatty acid cocktail and PUFAs solutions were supplemented at 1% v.v. in LB for overnight growth. MUFAs were first diluted 1:10 in ethanol and supplemented at 1% v.v. in LB for overnight growth. Undiluted vitamin E was supplemented from 0.1% to 10% in LB for overnight growth. MUFAs references: cis-vaccenic acid (VA; Cayman chemicals, 20023), palmitoleic acid (PA; Fisher Scientific, AC376910010), oleic acid (Sigma-Aldrich, 364525). PUFAs references: α-Linolenic Acid (ALA; Cayman chemicals, 90210), γ-Linolenic Acid (GLA; Cayman chemicals, 90220), dihomo-γ-Linolenic Acid (DGLA; Cayman chemicals, 90230). Antioxidants: Vitamin E (Sigma-Aldrich, T3251), N-acetyl-L-cystein (NAC; Sigma-Aldrich, A7250).

#### *C. elegans* FA measurements

Fatty acid measurements in bacteria and animals were performed as described before^68^. Briefly, lipids were extracted from whole animals using a 2:1 chloroform:methanol mixture and dried in liquid nitrogen. A calibrated phospholipid or triacylglycerol standard was added to the extracted lipids before separation by solid-phase exchange chromatography. Purified lipids were then converted into fatty acid methyl esters by incubation with methanol/2.5% H_2_SO_4_ before GC/MS analysis. All data are presented as percentage of the total fraction of fatty acids measured in the sample.

### Mammalian cell culture

MDA-MB-231 cells were obtained from the American Type Culture Collection (ATCC, Manassas, VA) and cultured in Dulbecco’s modified Eagle’s medium (Corning, 10-017-CV). T-47D cells were grown in RPMI-1640 media (Gibco, 11875119). All cell culture media were supplemented with 10% FBS (Hyclone, SH30910.03, lot AYG161519), 2 mM glutamine (Corning, 25-005-CI), and penicillin/streptomycin (Corning, 30-002-CI). Cells were cultured in a humidified incubator at 37°C with 5% CO_2_. In addition, cell lines were maintained at low passage numbers and regularly tested for mycoplasma.

### Mammalian drug sensitivity assays

Measurement of fractional viability (FV) and normalized growth rate inhibition (GR) were performed using the FLICK assay as previously described ^69^. Briefly, cells were seeded in 96-well optical-bottom black-walled plates (Greiner Bio-One, 655090) in 90 μL of media and allowed to adhere overnight. Density was optimized for each cell line depending on the growth rate of the cells and the duration of the assay. Dilutions of tamoxifen (Selleckchem, S1972) were prepared at 10x final concentration in media containing 50 μM SYTOX green (Thermo Fisher Scientific, S7020). 10 µL of drug combinations were supplemented to the cell culture media and fluorescence signals (excitation = 504 nm, emission = 523 nm) were recorded at the indicated time points on a Tecan Spark multimode plate reader. Total cell fluorescence was evaluated by lysing untreated cells at the start and experimental conditions at the end of the assay, using a supplement of 0.1% Triton X-100 (Fisher Scientific, BP151-500) diluted in PBS. Supplements: Ferrostatin-1 (Sigma-Aldrich, SML0583), ZVAD^70^.

### Gene expression profiling by RNA-seq

Total RNA was extracted using TRIzol (ThermoFisher), followed by DNase I (NEB) treatment and purified using the Direct-zol RNA mini-prep kit (Zymo research). RNA quality was verified by agarose gel electrophoresis. Briefly, multiplexed libraries were prepared using Cel-seq2^71^. Two to three biological replicates were sequenced with a NextSeq 500/550 High Output Kit v2.5 (75 Cycles) on a Nextseq500 sequencer. Paired end sequencing was performed; 13 cycles for read 1, six cycles for the illumina index and 73 cycles for read 2. Analysis was performed as previously detailed^72^, using a homemade DolphinNext pipeline^73^.

## Data availability

RNA-seq data have been deposited at GEO and is publicly available as of the date of publication under the following accession number: GSE186785. This study does not report original code.

## Acknowledgements

We thank members of the Walhout lab and Job Dekker for discussion and critical reading of the manuscript. We thank Leslie Shaw for providing the T-47D cell line. This work was supported by grants from the National Institutes of Health DK068429 and GM122502 to A.J.M.W., GM127559 to M.J.L., AG058950 to C.P.O. and GM122393 to A.G.G. Several bacterial and nematode strains used in this work were provided by the CGC, which is funded by the NIH Office of Research Infrastructure Programs (P40 OD010440).

## Author contributions

C.D., A.G.G. and A.J.M.W. conceived the study. C.D. and A.G.G. performed most experiments with technical help from M.W., H.D., O.P., Y.R. and A.F.V. performed the fatty acid measurements under supervision of C.P.O. H.N. engineered the *Δdhs-19* mutant strain. M.H. and C.D. performed the mammalian cell experiments under the supervision of M.L. C.D. and H.Z. performed and analyzed the RNA-seq experiments. The manuscript was written by C.D. and A.J.M.W. with editing by all other authors.

## Ethics declarations

The authors have no competing interests.

## Supplementary information

Supplementary information includes thirteen Supplementary Figures and six Supplementary Tables.

