## Supplemental Figures and Tables for "Bacterial diet modulates tamoxifen-induced death via host fatty acid metabolism"

Diot, et al. Supplementary Fig. 1

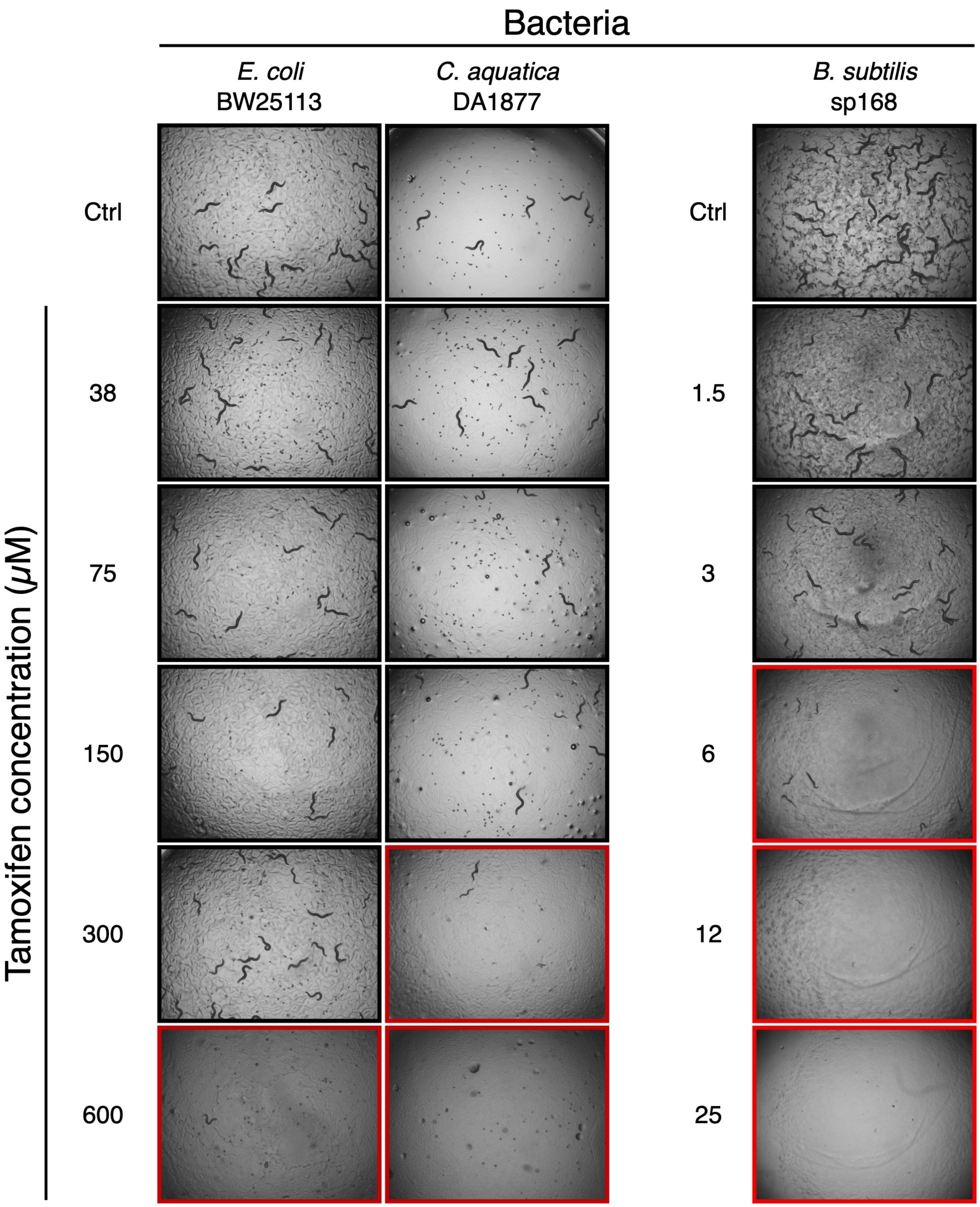

**Supplementary Fig. 1: tamoxifen toxicity in non-arrested L1 animals**

Bright-field images showing *C. elegans* supplemented with increasing doses of tamoxifen, fed *E. coli*, *C. aquatica*, and *B. subtilis*. In contrast to experiments presented in the main manuscript, experiments were conducted using *C. elegans* embryos rather than L1-arrested animals. Images were taken at 2x magnification after 48h exposure to tamoxifen. Representative of three independent experiments.

Diot, et al. Supplementary Fig. 2

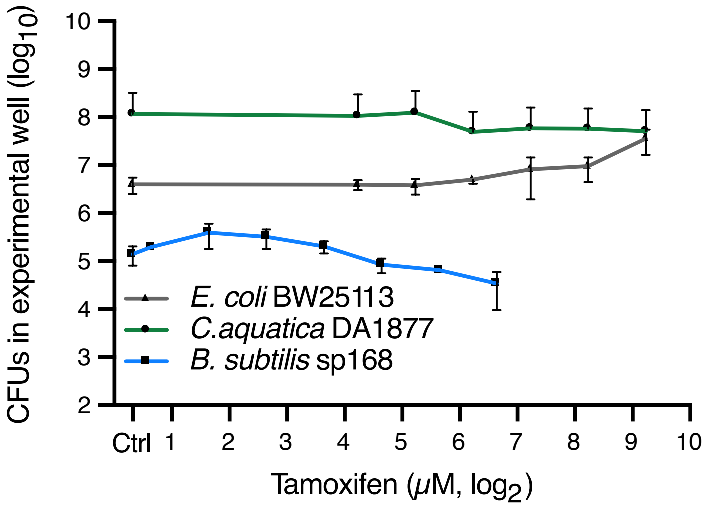

**Supplementary Fig. 2: assessment of bacterial viability on tamoxifen containing NGM plates**

Bacteria were seeded on tamoxifen containing NGM plates and incubated at room temperature for 48h. Bacterial lawns were recovered in M9 buffer, and colony-forming units were determined by plating serial dilutions on LB-agar plates. Mean of 2-3 biological replicates and SD are plotted.

*dhs-19* (VL1313)

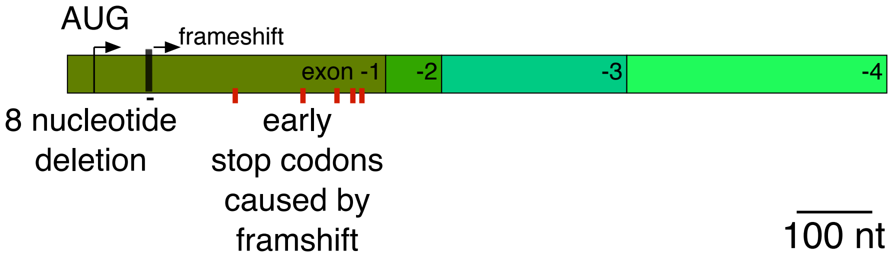

**Supplementary Fig. 3: Schematic of the *dhs-19* deletion**

To-scale cartoon of the exon 1 frameshift caused by an 8-nucleotide deletion in the *dhs-19* gene, in the engineered VL1313 strain.

Diot, et al. Supplementary Fig. 4

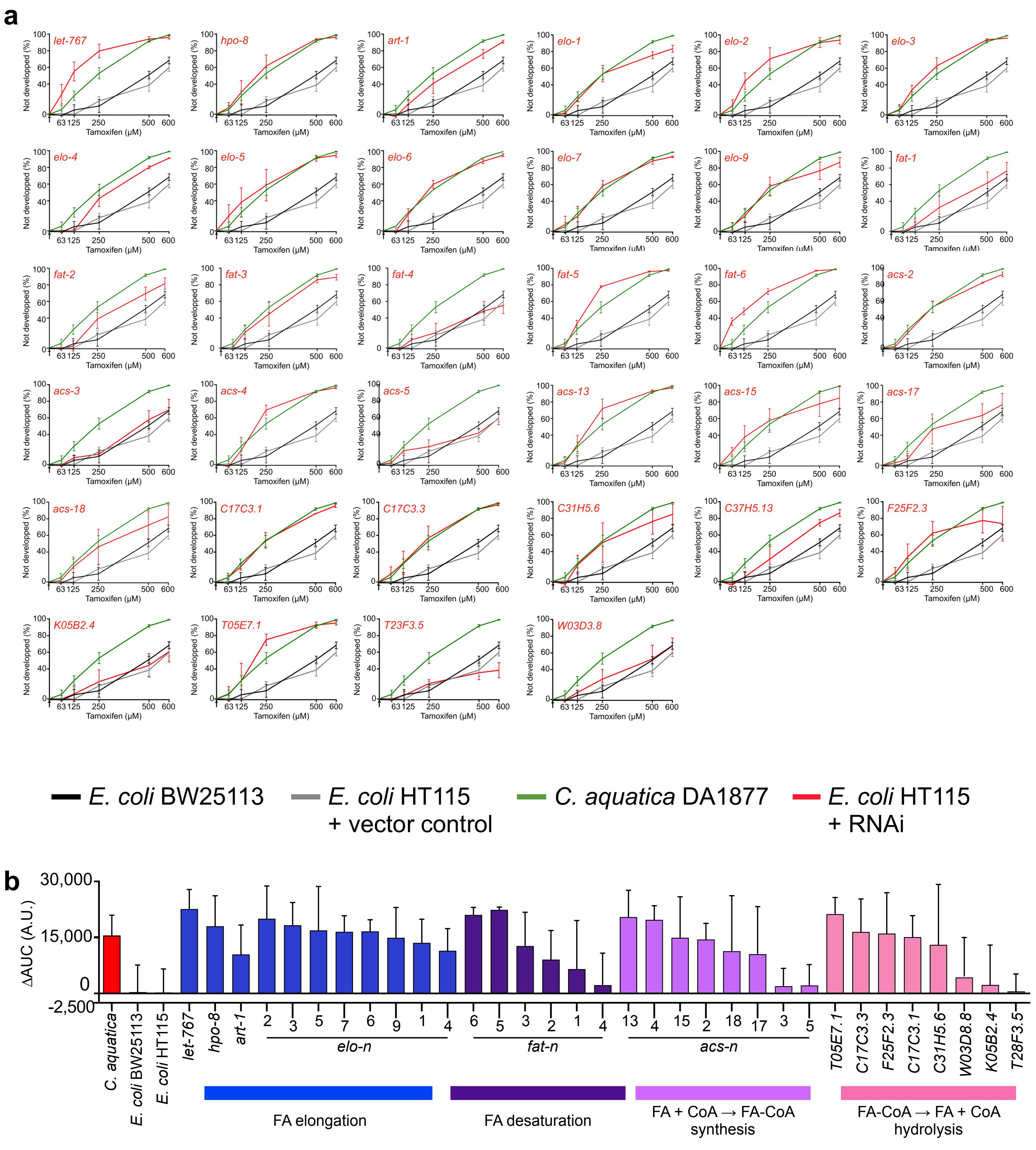

**Supplementary Fig. 4: Individual dose-response curves for animals exposed to RNAi of genes involved in the fatty acid biosynthesis.**

**a** All conditions were tested in parallel, and control conditions are plotted in each graph to facilitate the interpretation of the effect of individual RNAis. Data are represented as mean ±SEM of three biological replicates.

**b** AUCs presented in **Fig. 3e** with SD.

Diot, et al. Supplementary Fig. 5

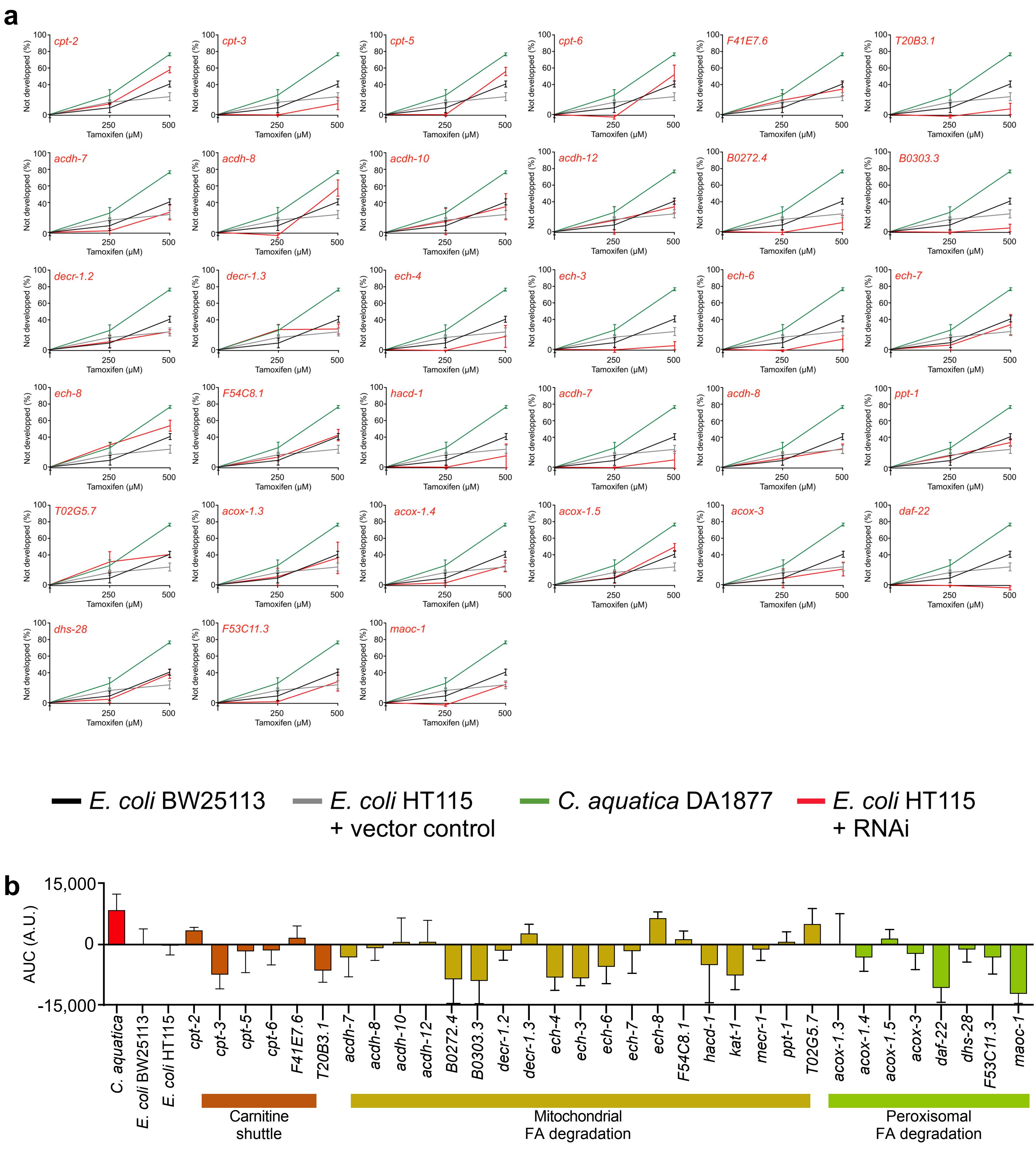

**Supplementary Fig. 5: Individual dose-response curves for animals exposed to RNAi of genes involved in the fatty acid degradation.**

**a** All conditions were tested in parallel, and control conditions are plotted in each graph to facilitate the interpretation of the effect of individual RNAis. Data are represented as mean ±SEM of three biological replicates.

**b** AUCs presented in **Fig. 3f** with SD.

Diot, et al. Supplementary Fig. 6

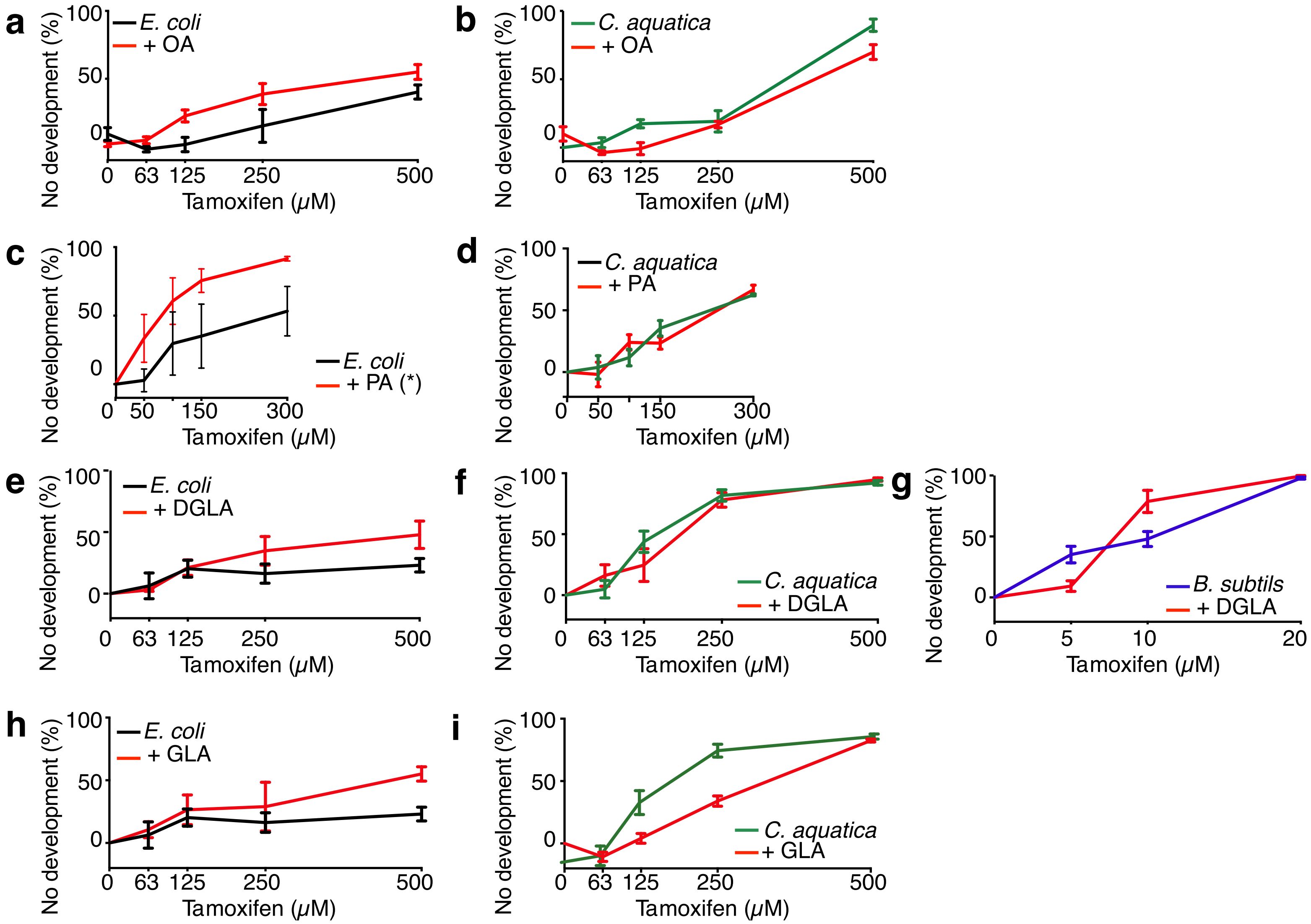

**Supplementary Fig. 6: DRCs of tamoxifen toxicity with a larger panel of FA supplements.**

**a**, **b** DRCs of tamoxifen toxicity with and without oleic acid supplementation on animals fed *E. coli* (**a**) or *C. aquatica* (**b**).

**c**, **d** DRCs of tamoxifen toxicity with and without palmitoleic acid supplementation on animals fed *E. coli* (**c**) or *C. aquatica* (**d**).

**e**-**g** DRCs of tamoxifen toxicity with and without di-homo-gamma-linoleic acid supplementation on animals fed *E. coli* (**e**), *C. aquatica* (**f**), or *B. subtilis* (**g**).

**h**, **i** DRCs of tamoxifen toxicity with and without gamma-linoleic acid supplementation on animals fed *E. coli* (**h**) or *C. aquatica* (**i**).

Diot, et al. Supplementary Fig. 7

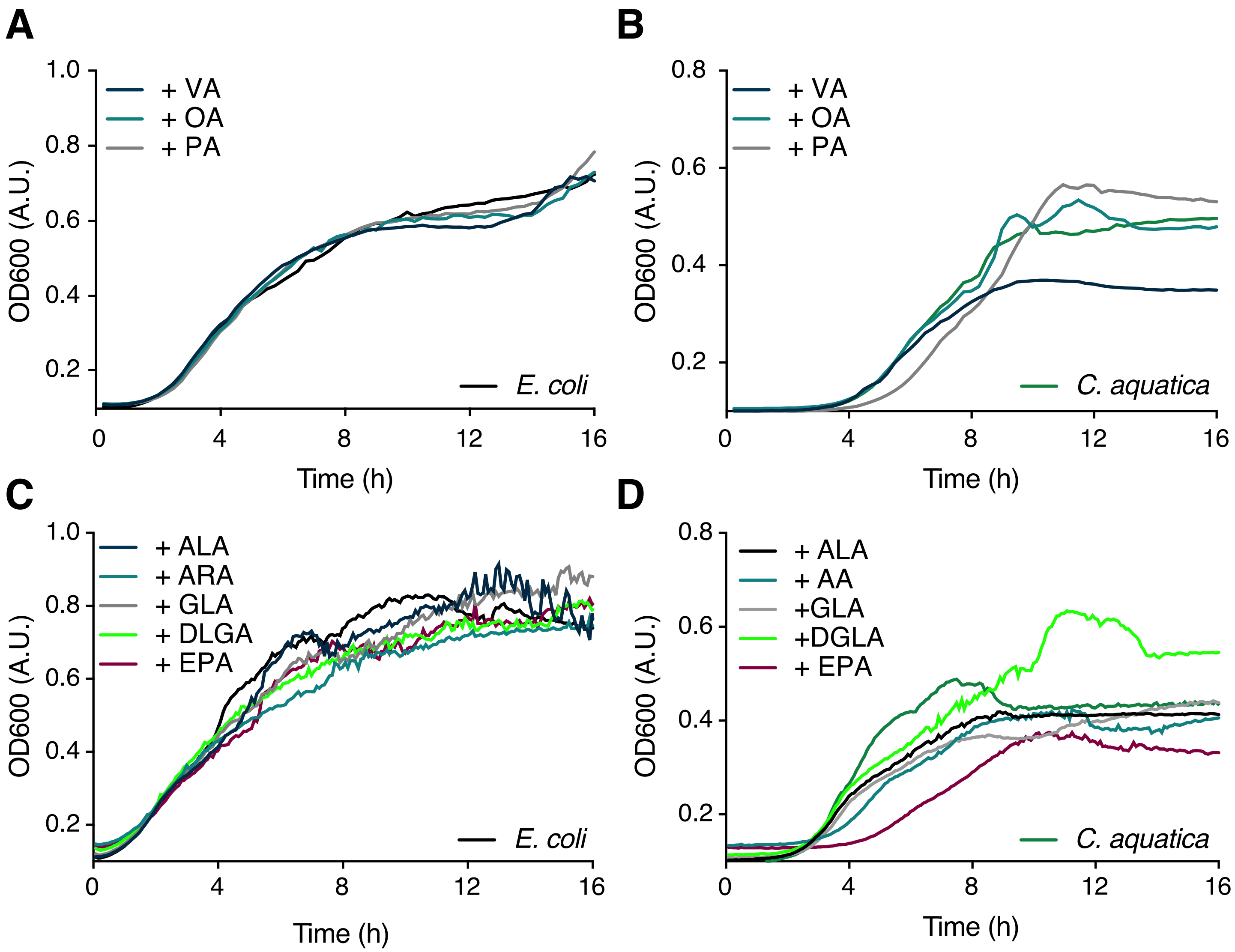

**Supplementary Fig. 7: Growth curves of *E. coli* and *C. aquatica* in presence of the FA supplements used in Fig. 4 and Supplementary Fig. S5.**

**a**, **c** Growth curves of *E. coli* supplemented with MUFAs (A) and PUFAs (C). For clarity, no error bars are reported in the graph, n=3.

**b**, **d** Growth curves of *C. aquatica* supplemented with MUFAs (A) and PUFAs (C). For clarity, no error bars are reported in the graph, n=3.

MUFA: mono-unsaturated FA, PUFA: poly-unsaturated FA, VA: cis-vaccenic acid, OA: oleic acid, PA: palmitoleic acid, ALA: alpha-linoleic acid, ARA: arachidonic acid, GLA: gamma-linoleic acid, DGLA: di-homo-gamma-linoleic acid, EPA: eicosapentaenoic acid.

Diot, et al. Supplementary Fig. 8

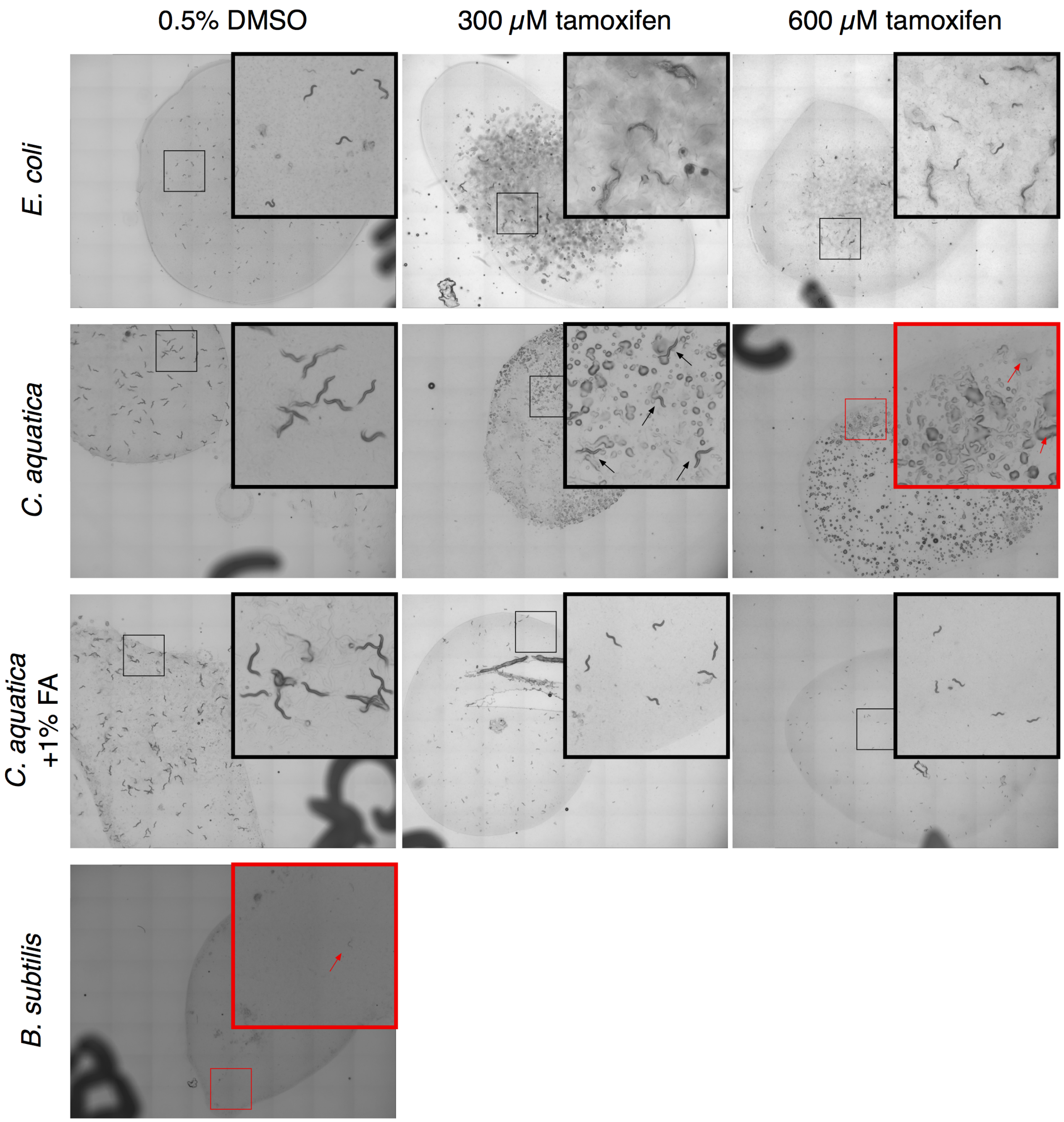

**Supplementary Fig. 8: tamoxifen toxicity in *C. elegans* fed metabolically inactive bacteria**

Bacterial powders were seeded on tamoxifen containing NGM plates, supplemented with antibiotics and devoid of peptone, and with FA cocktail when indicated. Dried plates were then seeded with L1-synchronized and pictures were taken at 72h. Images were taken at 2x magnification after 48h exposure to tamoxifen. Representative of three independent experiments.

### Diot, et al. Supplementary Fig. 9

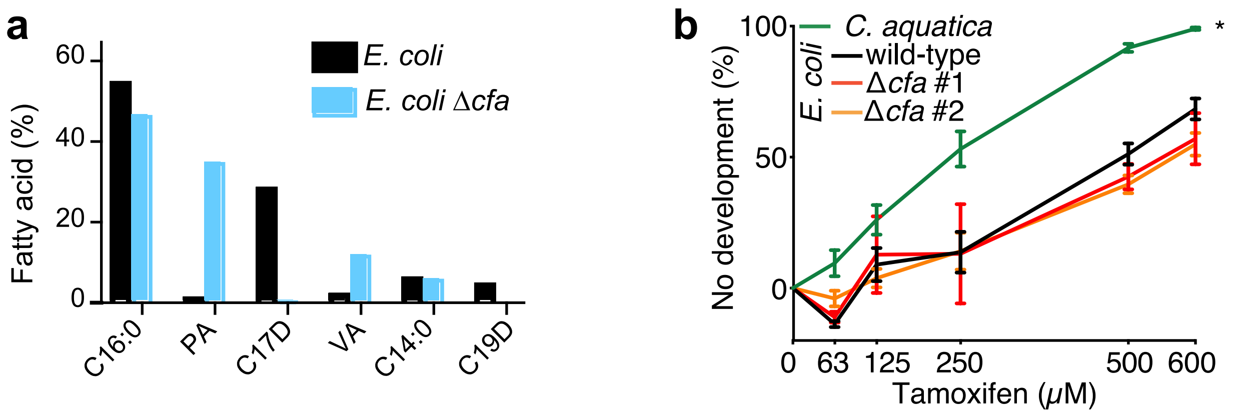

**Supplementary Fig. 9: tamoxifen toxicity in *C. elegans* fed in ∆*cfa* *E. coli* mutant**

**a** Fatty acid composition of *E. coli* BW25113 and ∆*cfa* mutant strains was assess by GC-MS.

**b** DRCs of tamoxifen toxicity on animals fed *C. aquatica*, *E. coli* BW25113, or *E. coli* ∆*cfa* mutant strains.

Diot, et al. Supplementary Fig. 10

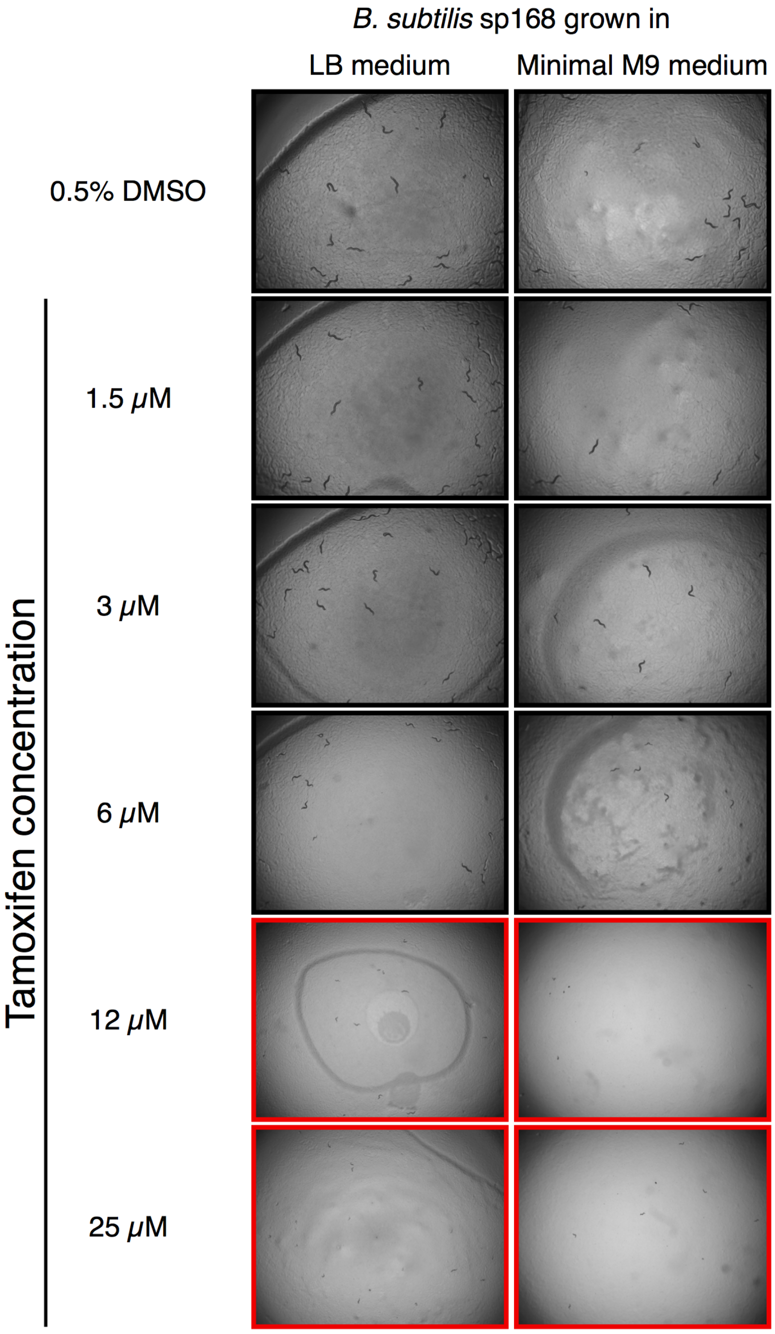

**Supplementary Fig. 10: tamoxifen toxicity in *C. elegans* fed *B. subtilis* grown in M9 minimal or LB media**

Bright-field images showing *C. elegans* supplemented with increasing doses of tamoxifen, fed *B. subtilis* grown in LB or M9 minimal media supplemented with glucose. Images were taken at 2x magnification after 48h exposure to tamoxifen.

Diot, et al. Supplementary Fig. 11

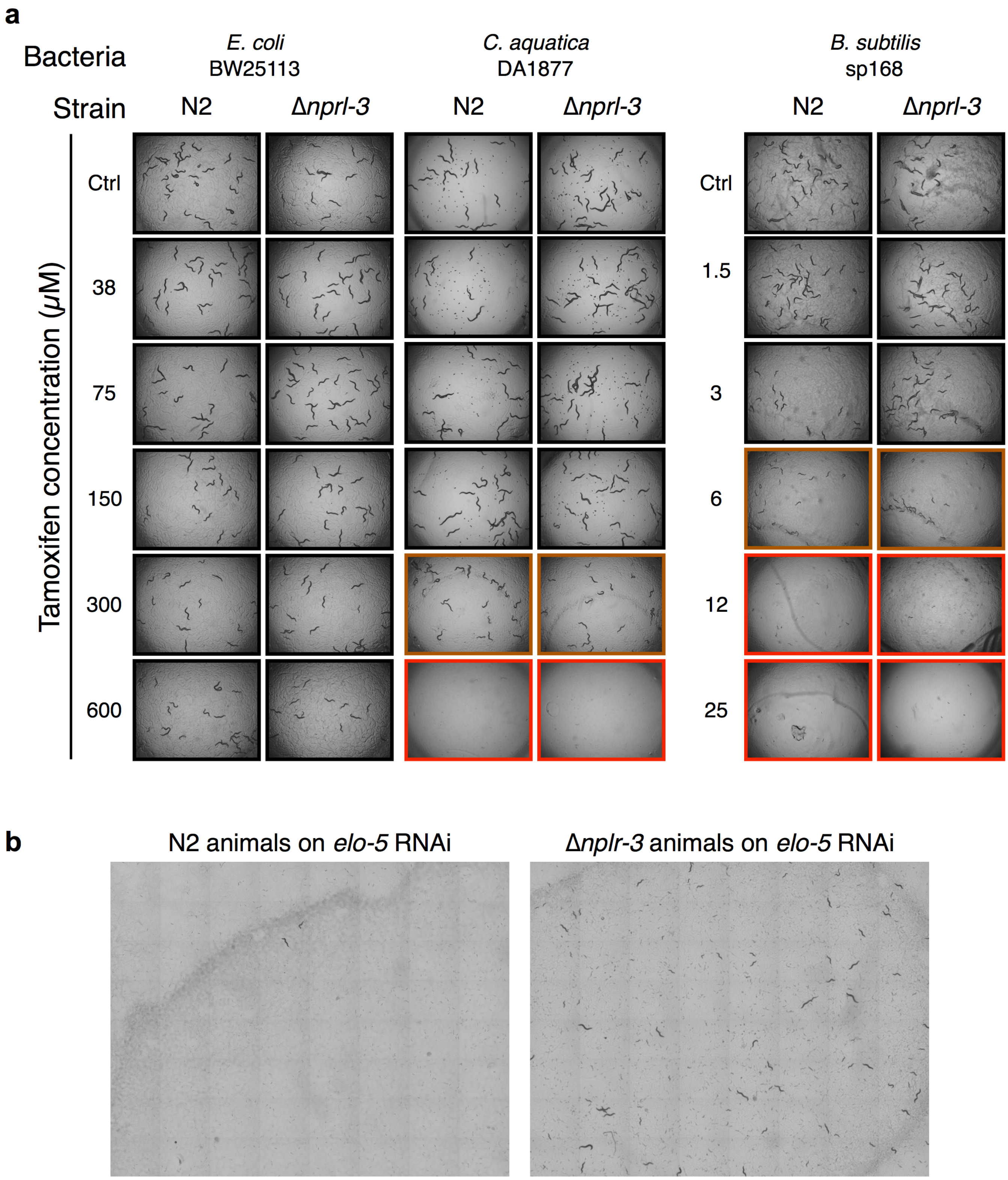

**Supplementary Fig. 11: tamoxifen toxicity in ∆*nprl-3* animals**

**a-b** experiments were conducted in parallel. Images are representative of three independent experiments.

**a** Bright-field images showing N2 and ∆*nprl-3* animal strains supplemented with increasing doses of tamoxifen, fed *E. coli, C. aquatica or B. subtilis.* Images were taken at 2x magnification after 48h exposure to tamoxifen.

**b** Bright-field images showing N2 and ∆*nprl-3* animal strains fed *E. coli* expressing double stranded RNA as indicated. Control indicates *E. coli* containing vector control plasmid (pL4440). F0 showed no phenotype. At 72h adult animals were removed from the plate and progeny was monitored daily. Images were taken at 10x magnification after 120h (48h post F0 removal).

Diot, et al. Supplementary Fig. 12

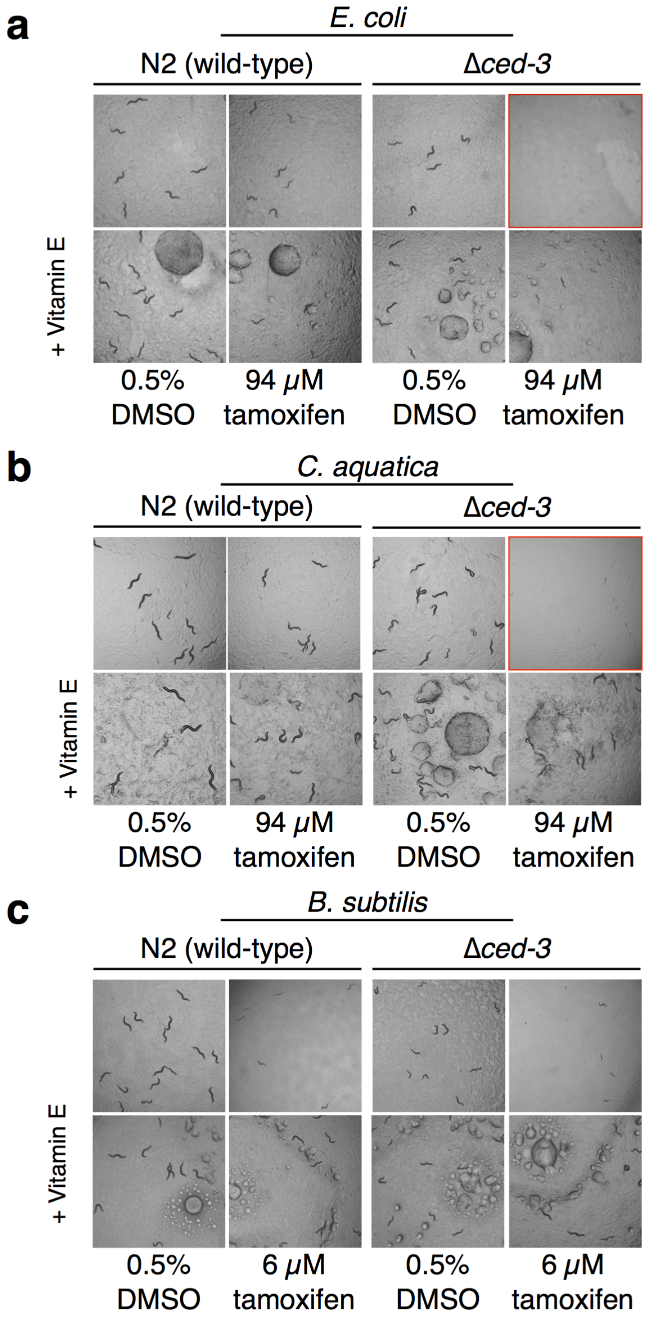

**Supplementary Fig. 12: diet-dependent potentiation of tamoxifen toxicity by a genetic deficiency to engage apoptosis**

**a** Bright-field images (left) of wild type and Δ*ced-3* animals fed *E. coli* supplemented or not with Vitamin E.

**b** Bright-field images (left) of wild type and Δ*ced-3* animals fed *C. aquatica* supplemented or not with Vitamin E.

**c** Bright-field images (left) of wild type and Δ*ced-3* animals fed *B. subtilis* supplemented or not with Vitamin E.

Diot, et al. Supplementary Fig. 13

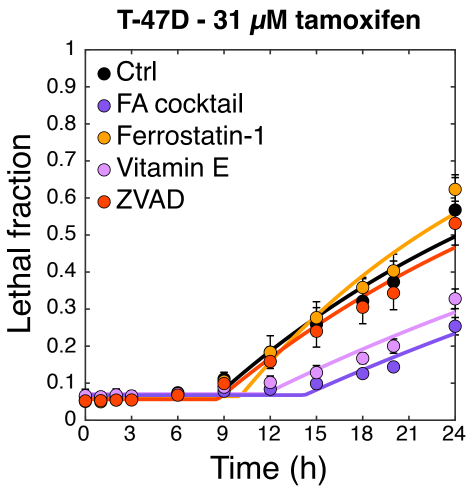

**Supplementary Fig. 13: effect of supplements on tamoxifen toxicity in ER-positive T-47D breast cancer cells**

Fractional viability of ER-positive T-47D breast cancer cells plotted as a function of time, in presence of the indicated supplements. Errors bars show SD, n=4.

Supplementary Table 1

Results of statistical analysis of tamoxifen dose response curves on animals knocked-down for genes involved in fatty acid bionsynthesis

| Control | Condition | p-value | Significance | Figure |
|---|---|---|---|---|
| *E. coli* | *C. aquatica* | <0.0001 | **** | Supplemental Figure 1 |
| *E. coli* | *let-727* | <0.0001 | **** | Supplemental Figure 1 |
| *E. coli* | *hpo-8* | <0.0001 | **** | Supplemental Figure 1 |
| *E. coli* | *art-1* | 0.0021 | ns | Supplemental Figure 1 |
| *E. coli* | *elo-1* | 0.0001 | *** | Supplemental Figure 1 |
| *E. coli* | *elo-2* | <0.0001 | **** | Supplemental Figure 1 |
| *E. coli* | *elo-3* | <0.0001 | **** | Supplemental Figure 1 |
| *E. coli* | *elo-4* | 0.0171 | ns | Supplemental Figure 1 |
| *E. coli* | *elo-5* | <0.0001 | **** | Supplemental Figure 1 |
| *E. coli* | *elo-6* | <0.0001 | **** | Supplemental Figure 1 |
| *E. coli* | *elo-7* | <0.0001 | **** | Supplemental Figure 1 |
| *E. coli* | *elo-9* | <0.0001 | **** | Supplemental Figure 1 |
| *E. coli* | *fat-1* | >0.9999 | ns | Supplemental Figure 1 |
| *E. coli* | *fat-2* | 0.311 | ns | Supplemental Figure 1 |
| *E. coli* | *fat-3* | <0.0001 | **** | Supplemental Figure 1 |
| *E. coli* | *fat-4* | >0.9999 | ns | Supplemental Figure 1 |
| *E. coli* | *fat-5* | <0.0001 | **** | Supplemental Figure 1 |
| *E. coli* | *fat-6* | <0.0001 | **** | Supplemental Figure 1 |
| *E. coli* | *acs-2* | <0.0001 | **** | Supplemental Figure 1 |
| *E. coli* | *acs-3* | >0.9999 | ns | Supplemental Figure 1 |
| *E. coli* | *acs-4* | <0.0001 | **** | Supplemental Figure 1 |
| *E. coli* | *acs-5* | >0.9999 | ns | Supplemental Figure 1 |
| *E. coli* | *acs-13* | <0.0001 | **** | Supplemental Figure 1 |
| *E. coli* | *acs-15* | <0.0001 | **** | Supplemental Figure 1 |
| *E. coli* | *acs-17* | 0.0522 | ns | Supplemental Figure 1 |
| *E. coli* | *acs-18* | 0.001 | ns | Supplemental Figure 1 |
| *E. coli* | *C17C3.1* | <0.0001 | **** | Supplemental Figure 1 |
| *E. coli* | *C17C3.3* | <0.0001 | **** | Supplemental Figure 1 |
| *E. coli* | *C31H5.6* | 0.0002 | *** | Supplemental Figure 1 |
| *E. coli* | *F25F2.3* | <0.0001 | **** | Supplemental Figure 1 |
| *E. coli* | *K05B2.4* | >0.9999 | ns | Supplemental Figure 1 |
| *E. coli* | *T05E7.1* | <0.0001 | **** | Supplemental Figure 1 |
| *E. coli* | *T28F3.5* | >0.9999 | ns | Supplemental Figure 1 |
| *E. coli* | *W03D8.8* | >0.9999 | ns | Supplemental Figure 1 |

Supplementary Table 2

Results of statistical analysis of tamoxifen dose response curves on animals knocked-down for genes involved in fatty acid degradation

| Control | Condition | p-value | Significance | Figure |
|---|---|---|---|---|
| *E. coli* | *C. aquatica* | 0.0004 | *** | Supplemental Figure 2 |
| *E. coli* | *cpt-2* | 0.9985 | ns | Supplemental Figure 2 |
| *E. coli* | *cpt-3* | 0.5142 | ns | Supplemental Figure 2 |
| *E. coli* | *cpt-5* | >0.9999 | ns | Supplemental Figure 2 |
| *E. coli* | *cpt-6* | >0.9999 | ns | Supplemental Figure 2 |
| *E. coli* | *F41E7.6* | 0.9999 | ns | Supplemental Figure 2 |
| *E. coli* | *T20B3.1* | 0.5586 | ns | Supplemental Figure 2 |
| *E. coli* | *acdh-7* | 0.9988 | ns | Supplemental Figure 2 |
| *E. coli* | *acdh-8* | 0.9997 | ns | Supplemental Figure 2 |
| *E. coli* | *acdh-10* | >0.9999 | ns | Supplemental Figure 2 |
| *E. coli* | *B0272.4* | 0.2354 | ns | Supplemental Figure 2 |
| *E. coli* | *B0303.3* | 0.1241 | ns | Supplemental Figure 2 |
| *E. coli* | *decr-1.2* | 0.9992 | ns | Supplemental Figure 2 |
| *E. coli* | *decr-1.3* | 0.9998 | ns | Supplemental Figure 2 |
| *E. coli* | *ech-4* | 0.4453 | ns | Supplemental Figure 2 |
| *E. coli* | *ech-3* | 0.1871 | ns | Supplemental Figure 2 |
| *E. coli* | *ech-6* | 0.8801 | ns | Supplemental Figure 2 |
| *E. coli* | *ech-7* | 0.9994 | ns | Supplemental Figure 2 |
| *E. coli* | *ech-8* | 0.9489 | ns | Supplemental Figure 2 |
| *E. coli* | *F54C8.1* | 0.9997 | ns | Supplemental Figure 2 |
| *E. coli* | *hacd-1* | 0.9423 | ns | Supplemental Figure 2 |
| *E. coli* | *kat-1* | 0.3731 | ns | Supplemental Figure 2 |
| *E. coli* | *mecr-1* | 0.9993 | ns | Supplemental Figure 2 |
| *E. coli* | *ppt-1* | >0.9999 | ns | Supplemental Figure 2 |
| *E. coli* | *T02G5.7* | 0.9987 | ns | Supplemental Figure 2 |
| *E. coli* | *acox-1.3* | 0.9999 | ns | Supplemental Figure 2 |
| *E. coli* | *acox-1.4* | 0.9986 | ns | Supplemental Figure 2 |
| *E. coli* | *acox-1.5* | 0.9995 | ns | Supplemental Figure 2 |
| *E. coli* | *acox-3* | 0.9988 | ns | Supplemental Figure 2 |
| *E. coli* | *daf-22* | 0.0141 | * | Supplemental Figure 2 |
| *E. coli* | *dhs-28* | 0.9997 | ns | Supplemental Figure 2 |
| *E. coli* | *F53C11.3* | 0.9988 | ns | Supplemental Figure 2 |
| *E. coli* | *maoc-1* | 0.0594 | ns | Supplemental Figure 2 |
| *E. coli* | *W03D8.8* | >0.9999 | ns | Supplemental Figure 1 |

Supplementary Table 3

Bacterial fatty acid composition. Results are expressed as the mean ± SD of three biological replicates.

| | | | *E. coli* | *E. coli* + 175 $\mu$M TAM | *C. aquatica* | *C. aquatica* + 175 $\mu$M TAM | *B. subtilis* |
|---|---|---|---|---|---|---|---|
| CPFA | | C17D | 31.84 ± 5.56 | 31.65 ± 1.86 | 3.78 ± 1.18 | 5.59 ± 2.26 | not detected |
| CPFA | | C19D | 13.85 ± 3.47 | 13.79 ± 0.92 | 0.84 ± 0.82 | 0.38 ± 0.37 | not detected |
| mmBCFA | | C15iso | not detected | 0.19 ± 0.19 | 0.01 ± 0.02 | 0.02 ± 0.02 | 15.52 ± 3.04 |
| mmBCFA | | C15ante-iso | not detected | not detected | not detected | not detected | 70.76 ± 2.66 |
| mmBCFA | | C17iso | 0.02 ± 0.04 | 0.02 ± 0.03 | 0.16 ± 0.28 | 0.06 ± 0.1 | 2.49 ± 0.46 |
| mmBCFA | | C17ante-iso | not detected | not detected | not detected | not detected | 7.35 ± 1.08 |
| MUFA | Palmitoleic Acid | C16:1n7 | 4.22 ± 3.77 | 4.41 ± 3.94 | 41.65 ± 9.06 | 44.54 ± 3.31 | 1.28 ± 0.13 |
| MUFA | Oleic Acid | C18:1n9 | 5.74 ± 3.18 | 5.58 ± 3.14 | 5.37 ± 1.16 | 8.62 ± 2.19 | not detected |
| MUFA | Vaccenic Acid | C18:1n7 | 0.9 ± 1.55 | 0.71 ± 1.24 | not detected | 0.13 ± 0.22 | not detected |
| SFA | Myristic Acid | C14:0 | 8.27 ± 0.57 | 8.05 ± 0.86 | 3.95 ± 0.83 | 4.00 ± 0.01 | 0.3 ± 0.01 |
| SFA | Palmitic Acid | C16:0 | 34.2 ± 6.74 | 34.93 ± 3.44 | 37.07 ± 6.29 | 36.29 ± 3.63 | 1.45 ± 0.27 |
| SFA | Stearic Acid | C18:0 | 0.55 ± 0.15 | 0.48 ± 0.14 | 0.68 ± 0.69 | 0.37 ± 0.36 | not detected |
| Total CPFA | | | 45.69 ± 8.66 | 45.44 ± 0.96 | 4.62 ± 1.97 | 5.97 ± 2.08 | not detected |
| Total mmBCFA | | | 0.02 ± 0.04 | 0.2 ± 0.22 | 0.18 ± 0.31 | 0.08 ± 0.12 | 96.57 ± 0.19 |
| Total MUFA | | | 10.86 ± 3.53 | 10.71 ± 4.17 | 47.02 ± 9.88 | 53.29 ± 2.48 | 1.28 ± 0.13 |
| Total PUFA | | | not detected | not detected | not detected | not detected | not detected |
| Total SFA | | | 43.02 ± 7.41 | 43.47 ± 3.25 | 41.69 ± 7.18 | 40.66 ± 3.51 | 2.11 ± 0.33 |

Supplementary Table 4

Fatty acid composition of *C. elegans*. Results are expressed as the mean ± SD of three biological replicates.

| Condition | | | *E. coli* BW25113 | *E. coli* BW25113 + 175 µM tamoxifen | *E. coli* HT115 + Control RNAi | *E. coli* HT115 + 175 µM tamoxifen + Control RNAi | *C. aquatica* | *C. aquatica* + 175 µM tamoxifen | *C. aquatica* + FA cocktail | *C. aquatica* + 175 µM tamoxifen + FA cocktail | *B. subtilis* |
|---|---|---|---|---|---|---|---|---|---|---|---|
| CPFA | C17D | | 07.43% ± 0.73 | 07.63% ± 0.43 | 04.73% ± 1.81 | 04.68% ± 2.03 | 00.89% ± 0.23 | 01.02% ± 0.33 | 00.53% ± 0.12 | 00.90% ± 0.24 | 0.79 ± 0.01 |
| CPFA | C19D | | 04.59% ± 0.18 | 04.85% ± 0.40 | 01.79% ± 0.92 | 01.92% ± 0.87 | 00.31% ± 0.06 | 00.17% ± 0.15 | 00.08% ± 0.08 | 00.12% ± 0.10 | not detected |
| mmBCFA | C15iso | | 02.13% ± 0.53 | 02.03% ± 0.49 | 01.41% ± 0.26 | 01.31% ± 0.28 | 01.56% ± 0.15 | 01.60% ± 0.25 | 01.07% ± 0.11 | 01.18% ± 0.08 | 2.01 ± 0.08 |
| mmBCFA | C15-anteiso | | not detected | not detected | not detected | not detected | not detected | not detected | not detected | not detected | 8.38 ± 0.24 |
| mmBCFA | C17iso | | 05.86% ± 0.46 | 06.44% ± 0.44 | 04.28% ± 1.42 | 04.68% ± 1.72 | 04.19% ± 0.40 | 04.42% ± 0.20 | 02.95% ± 0.23 | 02.87% ± 0.43 | 7.43 ± 0.07 |
| mmBCFA | C17-anteiso | | not detected | not detected | not detected | not detected | not detected | not detected | not detected | not detected | 16.93 ± 0.27 |
| MUFA | Palmitoleic Acid | C16:1n7 | 00.46% ± 0.05 | 00.38% ± 0.04 | 01.08% ± 0.21 | 00.84% ± 0.12 | 05.65% ± 0.96 | 05.06% ± 1.53 | 03.35% ± 1.37 | 03.48% ± 0.23 | 0.61 ± 0.01 |
| MUFA | Oleic Acid | C18:1n9 | 04.48% ± 1.42 | 03.74% ± 0.05 | 03.61% ± 1.40 | 03.55% ± 1.55 | 04.06% ± 0.06 | 04.04% ± 0.11 | 01.26% ± 0.44 | 01.36% ± 0.36 | 2.59 ± 0.26 |
| MUFA | Vaccenic Acid | C18:1n7 | 18.37% ± 1.32 | 17.16% ± 2.16 | 24.96% ± 8.64 | 24.78% ± 9.57 | 33.33% ± 1.60 | 34.16% ± 3.83 | 43.82% ± 0.94 | 45.14% ± 1.63 | 0.54 ± 0.07 |
| PUFA | γ-Linolenic acid | C18:3γ | 01.08% ± 0.94 | 01.13% ± 0.99 | 00.71% ± 0.30 | 00.60% ± 0.28 | 00.70% ± 0.61 | 00.74% ± 0.41 | 00.29% ± 0.08 | 00.42% ± 0.09 | 2.51 ± 0.19 |
| PUFA | Linoleic Acid | C18:2n6 | 10.25% ± 0.34 | 11.02% ± 0.23 | 09.82% ± 0.90 | 10.04% ± 1.69 | 09.90% ± 0.17 | 09.64% ± 0.14 | 04.22% ± 1.13 | 04.74% ± 1.35 | 4.65 ± 0.39 |
| PUFA | Arachadinoc Acid | C20:4n6 | 01.19% ± 0.02 | 01.33% ± 0.12 | 01.08% ± 0.38 | 01.15% ± 0.37 | 01.18% ± 0.27 | 01.15% ± 0.27 | 00.44% ± 0.09 | 00.55% ± 0.09 | 2.47 ± 0.06 |
| PUFA | Eicosapentaenoic acid | C20:5n3 | 23.32% ± 1.07 | 23.83% ± 0.88 | 28.87% ± 9.09 | 28.98% ± 8.33 | 21.41% ± 0.48 | 20.92% ± 1.38 | 23.97% ± 3.15 | 24.27% ± 1.14 | 29.26 ± 0.58 |
| PUFA | Dihomo-γ-linolenic acid | C20:3n6 | 03.84% ± 0.19 | 03.88% ± 0.31 | 02.75% ± 0.73 | 02.68% ± 0.81 | 02.67% ± 0.24 | 02.51% ± 0.39 | 02.00% ± 0.12 | 02.15% ± 0.16 | 6.53 ± 0.02 |
| PUFA | Eicosatetraenoic acid | C20:4n3 | 05.54% ± 0.12 | 05.38% ± 0.37 | 05.03% ± 1.07 | 04.94% ± 1.30 | 03.76% ± 0.20 | 03.55% ± 0.59 | 04.99% ± 0.47 | 04.47% ± 0.15 | 7.48 ± 0.24 |
| SFA | Myristic Acid | C14:0 | 0.27% ± 0.04 | 0.24% ± 0.04 | 00.19% ± 0.06 | 00.19% ± 0.08 | 00.11% ± 0.01 | 00.13% ± 0.02 | 00.09% ± 0.04 | 00.1% ± 0.01 | 0.31 ± 0.09 |
| SFA | Palmitic Acid | C16:0 | 03.68% ± 0.42 | 03.00% ± 0.62 | 03.27% ± 1.15 | 02.82% ± 0.67 | 03.17% ± 0.22 | 03.10% ± 0.51 | 03.05% ± 1.16 | 02.45% ± 0.20 | 1.48 ± 0.38 |
| SFA | Stearic Acid | C18:0 | 06.81% ± 0.80 | 07.18% ± 0.54 | 05.80% ± 1.47 | 06.50% ± 1.92 | 06.88% ± 1.24 | 07.56% ± 0.67 | 07.73% ± 1.08 | 05.59% ± 1.04 | 3.17 ± 0.37 |
| **Total CPFA** | | | 12.02 ± 0.85 | 12.48 ± 0.83 | 6.51 ± 2.72 | 6.6 ± 2.83 | 1.2 ± 0.17 | 1.19 ± 0.37 | 0.61 ± 0.2 | 1.02 ± 0.34 | 0.79 ± 0.01 |
| **Total mmBCFA** | | | 7.99 ± 0.97 | 8.47 ± 0.93 | 5.68 ± 1.5 | 5.98 ± 1.67 | 5.75 ± 0.51 | 6.03 ± 0.07 | 4.01 ± 0.31 | 4.05 ± 0.43 | 34.76 ± 0.31 |
| **Total MUFA** | | | 23.31 ± 0.18 | 21.28 ± 2.14 | 29.65 ± 8.77 | 29.16 ± 9.4 | 43.04 ± 2.05 | 43.27 ± 2.35 | 48.43 ± 0.94 | 49.98 ± 1.04 | 3.74 ± 0.32 |
| **Total PUFA** | | | 45.92 ± 1.12 | 47.35 ± 0.97 | 48.89 ± 8.44 | 48.74 ± 8.75 | 39.86 ± 1.09 | 38.72 ± 3.05 | 36.07 ± 2.46 | 36.81 ± 2.07 | 52.89 ± 0.85 |
| **Total SFA** | | | 10.75 ± 1.03 | 10.42 ± 0.39 | 9.26 ± 2.47 | 9.51 ± 2.44 | 10.16 ± 1.04 | 10.79 ± 0.87 | 10.87 ± 2.06 | 8.14 ± 0.93 | 7.82 ± 0.91 |

Supplementary Table 5. Page 1/2

Results of statistical analysis of fatty acid compositions between animals fed different diets ± tamoxifen ± FA cocktail

#### FA composition differences with E. coli BW25113

| | *E. coli* BW25113 + 175 µM TAM | *E. coli* HT115 + Control RNAi | *E. coli* HT115 + 175 µM TAM + Control RNAi | *C. aquatica* | *C. aquatica* + 175 µM TAM | *C. aquatica* + FA cocktail | *C. aquatica* + 175 µM TAM + FA cocktail |
|---|---|---|---|---|---|---|---|
| *E. coli* BW25113 | **no significant difference** | | | C17D ↓ 6.5% (**) (07.43% → 00.89%) | C17D ↓ 6.4% (**) (07.43% → 01.02%) | C17D ↓ 6.9% (**) (07.43% → 00.53%) | C17D ↓ 6.5% (**) (07.43% → 00.90%) |
| | | C19D ↓ 2.5% (*) (04.59% → 01.79%) | C19D ↓ 2.4%(*) (04.59% → 01.92%) | C19D ↓ 4.3% (***) (04.59% → 00.31%) | C19D ↓ 4.4% (***) (04.59% → 00.17%) | C19D ↓ 4.5% (**) (04.59% → 00.08%) | C19D ↓ 4.5% (***) (04.59% → 00.90%) |
| | | | | C17iso ↓ 1.7% (**) (05.86% → 01.56%) | | C17iso ↓ 2.9% (**) (05.86% → 01.07%) | C17iso ↓ 3.0% (**) (05.86% → 01.07%) |
| | | | | PA ↑ 5.0% (**) (00.46% → 05.65%) | PA ↑ 4.6% (*) (00.46% → 05.06%) | | PA ↑ 3.0% (**) (00.46% → 03.48%) |
| | | | | VA ↑ 15% (***) (18.37% → 33.33%) | VA ↑ 15.8% (*) (18.37% → 34.16%) | VA ↑ 25.5% (**) (18.37% → 43.82%) | VA ↑ 26.7% (**) (18.37% → 45.14%) |
| | | | | | | LA ↓ 6% (*) (10.25% → 04.22%) | LA ↓ 5.5% (*) (10.25% → 04.74%) |
| | | | | C20:5n3 ↓ 1.9% (*) (23.32% → 21.41%) | | | |
| | | | | DGLA ↓ 1.2% (*) (03.84% → 02.67%) | DGLA ↓ 1.3% (*) (03.84% → 02.51%) | DGLA ↓ 1.8% (**) (03.84% → 02.00%) | DGLA ↓ 1.7% (**) (03.84% → 02.15%) |
| | | | | ETA ↓ 1.8% (**) (05.54% → 03.76%) | ETA ↓ 2.0% (*) (05.54% → 03.55%) | ETA ↓ 2.0% (*) (05.54% → 04.99%) | ETA ↓ 1.1% (**) (05.54% → 04.47%) |

#### FA composition differences with E. coli HT115

| | *E. coli* BW25113 | *E. coli* BW25113 + 175 µM TAM | *E. coli* HT115 + 175 µM TAM + Control RNAi | *C. aquatica* | *C. aquatica* + 175 µM TAM | *C. aquatica* + FA cocktail | *C. aquatica* + 175 µM TAM + FA cocktail |
|---|---|---|---|---|---|---|---|
| *E. coli* HT115 | | | **no significant difference** | C17D ↓ 4.7% (*) (04.73% → 00.89%) | C17D ↓ 4.3% (*) (04.73% → 01.02%) | C17D ↓ 4.8% (*) (04.73% → 00.53%) | C17D -4.5% (*) (04.73% → 00.90%) |
| | C19D ↑ 2.5% (*) (01.79% → 04.59%) | C19D ↑ 2.5% (*) 01.79% → 04.85% | | PA ↑ 4.5% (*) (01.08% → 05.65%) | | | PA +2.3% (**) (01.08% → 03.48%) |
| | | | | VA ↑ 11.8% (24.96% → 33.33%) | VA ↑ 12.6% (*) (24.96% → 34.16%) | VA ↑ 22.3% (*) (24.96% → 43.82%) | VA +23.6% (*) (24.96% → 45.14%) |
| | | | | | | LA ↓ 5.8% (**) (09.82% → 04.22%) | LA -5.4% (**) (09.82% → 04.74%) |

#### FA composition differences with E. coli BW25113 + 175 µM TAM

| | *E. coli* BW25113 | *E. coli* HT115 + Control RNAi | *E. coli* HT115 + 175 µM TAM + Control RNAi | *C. aquatica* | *C. aquatica* + 175 µM TAM | *C. aquatica* + FA cocktail | *C. aquatica* + 175 µM TAM + FA cocktail |
|---|---|---|---|---|---|---|---|
| *E. coli* BW25113 + 175 µM TAM | **no significant difference** | | | C17D ↓ 6.7% (**) (07.63% → 00.89%) | C17D ↓ 6.6% (**) (07.63% → 01.02%) | C17D ↓ 7.1% (***) (07.63% → 00.53%) | C17D ↓ 6.7% (***) (07.63% → 00.90%) |
| | | C19D ↓ 2.5% (*) (04.85% → 01.79%) | C19D ↓ 2.6% (**) (04.85% → 01.92%) | C19D ↓ 4.5% (**) (04.85% → 00.31%) | C19D ↓ 4.7% (***) (04.85% → 00.17%) | C19D ↓ 4.8% (**) (04.85% → 00.08%) | C19D ↓ 4.7% (**) (04.85% → 00.12%) |
| | | | | C17iso ↓ 2.2% (**) (06.44% → 04.19%) | C17iso ↓ 2.0% (*) (06.44% → 04.42%) | C17iso ↓ 3.4% (**) (06.44% → 02.95%) | C17iso ↓ 3.6% (**) (06.44% → 02.87%) |
| | | | | PA ↑ 5.2% (*) (00.38% → 05.65%) | PA ↑ 4.7% (*) (00.38% → 05.06%) | PA ↓ 2.5% (*) (00.38% → 03.35%) | PA ↓ 3.1% (**) (00.38% → 03.48%) |
| | | | | | | | OA ↓ 2.4% (**) (03.74% → 01.36%) |
| | | | | VA ↑ 16.2% (***) (17.16% → 33.33%) | VA ↑ 17.0% (**) (17.16% → 34.16%) | VA ↑ 26.7% (**) (17.16% → 43.82%) | VA ↑ 28.0% (**) (17.16% → 45.14%) |
| | | | | LA ↓ 1.1% (*) (11.02% → 09.90%) | LA ↓ 1.4% (*) (11.02% → 09.64%) | LA ↓ 6.8% (*) (11.02% → 04.22%) | LA ↓ 6.2% (*) (11.02% → 04.74%) |
| | | | | C20:5n3 ↓ 2.4% (*) (23.83% → 21.41%) | C20:5n3 ↓ 2.9% (*) (23.83% → 20.92%) | | |
| | | | | DGLA ↓ 1.2% (*) (03.88% → 02.67%) | DGLA ↓ 1.3% (**) (03.88% → 02.51%) | DGLA ↓ 1.9% (**) (03.88% → 02.00%) | DGLA ↓ 1.7% (**) (03.88% → 02.15%) |
| | | | | ETA ↓ 1.6% (*) (05.38% → 03.76%) | ETA ↓ 1.8% (**) (05.38% → 03.55%) | | |

#### FA composition differences with C. aquatica + 175 µM + FA cocktail

| | *E. coli* BW25113 | *E. coli* BW25113 + 175 µM TAM | *E. coli* HT115 + Control RNAi | *E. coli* HT115 + 175 µM TAM + Control RNAi | *C. aquatica* | *C. aquatica* + 175 µM TAM | *C. aquatica* + FA cocktail |
|---|---|---|---|---|---|---|---|
| *C. aquatica* + 175 µM + FA cocktail | C17D ↑ 6.5% (**) (00.90% → 07.43%) | C17D ↑ 6.7% (***) (00.90% → 07.63%) | C17D ↑ 4.5% (*) (00.90% → 04.73%) | C17D ↑ 4.5% (*) (00.90% → 04.68%) | C17iso ↑ 1.3% (*) (02.87% → 04.19%) | C17iso ↑ 1.6% (*) (02.87% → 04.42%) | **no significant difference** |
| | C19D ↑ 4.5% (***) (00.90% → 04.59%) | C19D ↑ 4.7% (**) (00.12% → 04.85%) | | C19D ↑ 2.1% (*) (00.12% → 01.92%) | | | |
| | C17iso ↑ 3.0% (**) (01.07% → 05.86%) | C17iso ↑ 3.6% (**) (02.87% → 06.44%) | | | | | |
| | PA ↓ 3.0% (**) (03.48% → 00.46%) | PA ↑ 3.1% (**) (03.48% → 00.38%) | PA ↓ 2.3% (**) (03.48% → 01.08%) | PA ↓ 2.6% (**) (03.48% → 00.84%) | | | |
| | | OA ↑ 2.4% (**) (01.36% → 03.74%) | | | PA ↑ 2.1% (*) (03.48% → 05.65%) | OA ↑ 2.7% (**) (01.36% → 04.04%) | |
| | VA ↓ 26.7% (**) (45.14% → 18.37%) | VA ↓ 28.0% (**) (45.14% → 17.16%) | VA ↓ 23.6% (*) (45.14% → 24.96%) | VA ↓ 23.9% (*) (45.14% → 24.78%) | OA ↑ 2.7% (**) (01.36% → 04.06%) | VA ↓ 11% (*) (45.14% → 34.16%) | |
| | LA ↑ 5.5% (*) (04.74% → 0.25% ) | LA ↑ 6.2% (*) (04.74% → 11.02%) | LA ↑ 5.4% (**) (04.74% → 09.82%) | LA ↑ 6.0% (*) (04.74% → 10.04%) | LA ↑ 5.2% (*) (04.74% → 09.90%) | LA ↑ 4.9% (*) (04.74% → 09.64%) | |
| | DGLA ↑ 1.7% (**) (02.15% → 03.84%) | DGLA ↑ 1.7% (**) (02.15% → 03.88%)) | | | | | |
| | ETA ↑ 1.1% (**) (04.47% → 05.54%) | | | | | | |

Supplementary Table 5. Page 2/2

Results of statistical analysis of fatty acid compositions between animals fed different diets ± tamoxifen ± FA cocktail

##### FA composition differences with C. aquatica

| | *E. coli* BW25113 | *E. coli* BW25113 + 175 μM TAM | *E. coli* HT115 + Control RNAi | *E. coli* HT115 + 175 μM TAM + Control RNAi | *C. aquatica* + 175 μM TAM | *C. aquatica* + FA cocktail | *C. aquatica* + 175 μM TAM + FA cocktail |
|---|---|---|---|---|---|---|---|
| *C. aquatica* | C17D ↑ 6.5% (**) (00.89% → 07.43%) | C17D ↑ 6.7% (**) (00.89% → 07.63%) | C17D ↑ 4.7% (*) (00.89% → 04.73%) | C17D ↑ 4.5% (*) (0.89%) → 4.68% ) | **no significant difference** | | |
| | C19D ↑ 4.3% (***) (04.59% → 00.31%) | C19D ↑ 4.5% (**) (00.31% → 04.85%) | | | | | |
| | C17iso ↑ 1.7% (**) (04.59% → 00.31%) | C17iso ↑ 2.2% (**) (00.31% → 04.85%) | | | | C17iso ↓ 1.2% (**) (04.19% → 02.95%) | C17iso ↓ 1.3% (*) (04.19% → 02.87%) |
| | PA ↓ 5.0% (**) (05.65% → 00.46%) | PA ↓ 5.2% (*) (00.31% → 04.85%) | PA ↓ 4.5% (*) (05.65% → 01.08%) | PA ↓ 4.7% (*) (05.65% → 00.84%) | | PA ↓ 2.3% (*) (05.65% → 03.35%) | PA ↓ 2.1% (*) (05.65% → 03.48%) |
| | | | | | | | OA ↓ 2.7% (**) (04.06% → 01.36%) |
| | VA ↓ 15% (***) (33.33% → 18.37%) | VA ↓ 16.2% (***) (33.33% → 17.16%) | VA ↓ 11.8% ((33.33% → 24.96%) | VA ↓ 12.1% (33.33% → 24.78%) | | VA ↑ 10.5% (**) (33.33% → 43.82%) | VA ↑ 11.8% (*) (33.33% → 45.14%) |
| | | LA ↑ 1.1% (*) (33.33% → 17.16%) | | | | LA ↓ 5.7% (*) (09.90% → 04.22%) | LA ↓ 5.2% (*) (09.90% → 04.74%) |
| | C20:5n3 ↑ 1.9% (*) (23.32% → 21.41%) | C20:5n3 ↑ 2.4% (*) (21.41% → 23.83%) | | | | | |
| | DGLA ↑ 1.2% (*) (02.67% → 03.84%) | DGLA ↑ 1.2% (*) (02.67% → 03.88%) | | | | | |
| | ETA ↑ 1.8% (**) (05.54% → 03.76%) | ETA ↑ 1.6% (*) (03.76% → 05.38%) | | | | ETA ↑ 1.2% (*) (03.76% → 04.99%) | |

##### FA composition differences with E. coli HT115 + 175 μM TAM

| | *E. coli* BW25113 | *E. coli* BW25113 + 175 μM TAM | *E. coli* HT115 + Control RNAi | *C. aquatica* | *C. aquatica* + 175 μM TAM | *C. aquatica* + FA cocktail | *C. aquatica* + 175 μM TAM + FA cocktail |
|---|---|---|---|---|---|---|---|
| *E. coli* HT115 + 175 μM TAM | | | **no significant difference** | C17D -4.5% (*) (04.68% → 00.89%)" | C17D -4.4% (*) (04.68% → 01.02%) | C17D -4.9% (*) (04.68% → 00.53%) | C17D -4.5% (*) (04.68% → 00.90%) |
| | C19D ↑ 2.4%(*) (01.92% → 04.59%) | C19D ↑ 2.6% (**) (01.92% → 04.85%) | | | C19D 2.0% (*) (01.92% → 00.17%) | C19D -2.1% (*) (01.92% → 00.08%) | C19D -2.1% (*) (01.92% → 00.12%) |
| | | | | PA +4.7% (*) (00.84% → 05.65%) | PA +4.2% (*) (00.84% → 05.06%) | | PA +2.6% (**) (00.84% → 03.48%) |
| | | | | VA +12.1% (24.78% → 33.33%) | VA +12.9% (*) (24.78% → 34.16%) | VA +22.6% (*) (24.78% → 43.82%) | VA +23.9% (*) (24.78% → 45.14%) |
| | | | | | | LA -6.5% (**) (10.04% → 04.22%) | LA -6.0% (*) (10.04% → 04.74%) |

##### FA composition differences with C. aquatica + 175 μM

| | *E. coli* BW25113 | *E. coli* BW25113 + 175 μM TAM | *E. coli* HT115 + Control RNAi | *E. coli* HT115 + 175 μM TAM + Control RNAi | *C. aquatica* | *C. aquatica* + FA cocktail | *C. aquatica* + 175 μM TAM + FA cocktail |
|---|---|---|---|---|---|---|---|
| *C. aquatica* + 175 μM | C17D ↑ 6.4% (**) (01.02% → 07.43%) | C17D ↑ 6.6% (**) (01.02% → 07.63%) | C17D ↑ 4.3% (*) (01.02% → 04.73%) | C17D ↑ 4.4% (*) (01.02% → 04.68%) | **no significant difference** | | |
| | C19D ↑ 4.4% (***) ((01.02% → 07.43%) | C19D ↑ 4.7% (***) (00.17% → 04.85%) | | C19D ↑ 2.0% (*) (00.17% → 01.92%) | | | |
| | | C17iso ↑ 2.0% (*) (04.42% → 06.44%) | | | | C17iso ↓ 1.5% (*) (04.42% → 02.95%) | C17iso ↓ 1.6% (*) (04.42% → 02.87%) |
| | PA ↓ 4.6% (*) (05.06% → 00.46%) | PA ↓ 4.7% (*) (05.06% → 00.38%) | | PA ↓ 4.2% (*) (05.06% → 00.84%) | | | |
| | | | | | | OA ↓ 2.8% (*) (04.04% → 01.26%) | OA ↓ 2.7% (**) (04.04% → 01.36%) |
| | VA ↓ 15.8% (*) (34.16% → 18.37%) | VA ↓ 17.0% (**) (34.16% → 17.16%) | VA ↓ 12.6% (*) (34.16% → 24.96%) | VA ↓ 12.9% (*) (34.16% → 24.78%) | | VA ↑ 9.7% (*) (34.16% → 43.82%) | VA ↑ 11% (*) (34.16% → 45.14%) |
| | | LA ↑ 1.4% (*) (09.64% → 11.02%) | | | | LA ↓ 5.4% (*) (09.64% → 04.22%) | LA ↓ 4.9% (*) (09.64% → 04.74%) |
| | DGLA ↑ 1.3% (*) (02.51% → 03.84%) | C20:5n3 ↑ 2.9% (*) (20.92% → 23.83%) | | | | | |
| | ETA ↑ 2.0% (*) (03.55% → 05.54%) | DGLA ↑ 1.3% (**) (02.51% → 03.88%) | | | | | |
| | ETA ↑ 1.8% (**) (03.76% → 05.54%) | ETA ↑ 1.8% (**) (03.55% → 05.38%) | | | | ETA ↑ 1.4% (*) (03.55% → 04.99%) | |

##### FA composition differences with C. aquatica + FA cocktail

| | *E. coli* BW25113 | *E. coli* BW25113 + 175 μM TAM | *E. coli* HT115 + Control RNAi | *E. coli* HT115 + 175 μM TAM + Control RNAi | *C. aquatica* | *C. aquatica* + 175 μM TAM | *C. aquatica* + 175 μM TAM + FA cocktail |
|---|---|---|---|---|---|---|---|
| *C. aquatica* + FA cocktail | C17D ↑ 6.9% (**) (00.53% → 07.43%) | C17D ↑ 7.1% (***) (00.53% → 07.63%) | C17D ↑ 4.8% (*) (00.53% → 04.73%) | C17D ↑ 4.9% (*) (00.53% → 04.68%) | | | **no significant difference** |
| | C19D ↑ 4.5% (**) (00.08% → 04.59%) | C19D ↑ 4.8% (**) (00.53% → 07.63%) | | C19D ↑ 2.1% (*) (00.08% → 01.92%) | | | |
| | C17iso ↑ 2.9% (**) (01.07% → 05.86%)) | C17iso ↑ 3.4% (**) (02.95% → 06.44%) | | | C17iso ↑ 1.2% (**) (02.95% → 04.19%) | C17iso ↑ 1.5% (*) (02.95% → 04.42%) | |
| | | PA ↓ 2.5% (*) (03.35% → 00.38%) | | | PA ↑ 2.3% (*) (03.35% → 05.65%) | | |
| | | | | | | OA ↑ 2.8% (*) (01.26% → 04.04%) | |
| | VA ↓ 25.5% (**) (43.82% → 18.37%) | VA ↓ 26.7% (**) (43.82% → 17.16%) | VA ↓ 22.3% (*) (00.53% → 04.73%) | VA ↓ 22.6% (*) (43.82% → 24.78%) | VA ↓ 10.5% (**) (43.82% → 33.33%) | VA ↓ 9.7% (*) (01.26% → 04.04%) | |
| | LA ↑ 6% (*) (04.22% → 0.25% ) | LA ↑ 6.8% (*) (04.22% → 11.02%) | LA ↑ 5.8% (**) (00.53% → 04.73%) | LA ↑ 6.5% (**) (04.22% → 10.04%) | LA ↑ 5.7% (*) (04.22% → 09.90%) | LA ↑ 5.4% (*) (04.22% → 09.64%) | |
| | DGLA ↑ 1.8% (**) (02.00% → 03.84%) | DGLA ↑ 1.9% (**) (02.00% → 03.88%) | | | | | |
| | ETA ↑ 2.0% (*) (04.99% → 05.54%) | | | | ETA ↓ 1.2% (*) (04.99% → 03.76%) | ETA ↓ 1.4% (*) (04.99% → 03.55%) | |
